# Sex-specific expression of the human NAFLD-NASH transcriptional signatures in the liver of medaka with a history of ancestral bisphenol A exposure

**DOI:** 10.1101/2024.05.19.594843

**Authors:** Sourav Chakraborty, Santosh Anand, Ramji Kumar Bhandari

## Abstract

The progression of fatty liver disease to non-alcoholic steatohepatitis (NASH) is a leading cause of death in humans. Lifestyles and environmental chemical exposures can increase the susceptibility of humans to NASH. In humans, the presence of bisphenol A (BPA) in urine is associated with fatty liver disease, but whether ancestral BPA exposure leads to the activation of human NAFLD-NASH-associated genes in the unexposed descendants is unclear. In this study, using medaka fish as an animal model for human NAFLD, we investigated the transcriptional signatures of human NAFLD-NASH and their associated roles in the pathogenesis of the liver of fish who were not directly exposed but their ancestors were exposed to BPA during embryonic and perinatal development three generations prior. Comparison of bulk RNA-Seq data of the liver in BPA lineage male and female medaka with publicly available human NAFLD-NASH patient data revealed transgenerational alterations in the transcriptional signature of human NAFLD-NASH in medaka liver. Twenty percent of differentially expressed genes (DEGs) were upregulated in both human NAFLD patients and medaka. Specifically in females, among the total shared DEGs in the liver of BPA lineage fish and NAFLD patient groups, 27.69% DEGs were downregulated and 20% DEGs were upregulated. Off all DEGs, 52.31% DEGs were found in ancestral BPA-lineage females, suggesting that NAFLD in females shared majority of human NAFLD gene networks. Pathway analysis revealed beta-oxidation, lipoprotein metabolism, and HDL/LDL-mediated transport processes linked to downregulated DEGs in BPA lineage males and females. In contrast, the expression of genes encoding lipogenesis-related proteins was significantly elevated in the liver of BPA lineage females only. BPA lineage females exhibiting activation *of myc, atf4, xbp1*, *stat4*, and cancerous pathways, as well as inactivation of *igf1*, suggest their possible association with an advanced NAFLD phenotype. The present results suggest that gene networks involved in the progression of human NAFLD and the transgenerational NAFLD in medaka are conserved and that medaka can be an excellent animal model to understand the development and progression of liver disease and environmental influences in the liver.

## Introduction

Nonalcoholic fatty liver disease (NAFLD) is a leading liver disease in Western countries, and its incidence is on the rise worldwide^1^. According to the “two-hit hypothesis,” steatosis or NAFLD is caused by more than 5% accumulation of fat in hepatocytes in the first hit^2^. Consequently, excess fat accumulation causes the second hit, which involves oxidative stress, pro-inflammatory cytokines synthesis, and ultimately, apoptosis^3^. Underdiagnosed and untreated NAFLD can lead to necro-inflammatory changes in the liver, which is known as non-alcoholic steatohepatitis (NASH)^4^. In NASH, liver stellate cells (HSCs) release collagen type I and III and fibronectin, which promote fibrosis through chronic liver inflammation^5^. As the disease progresses, ballooning degeneration, lobular inflammation, collagen fiber deposition, and cell death eventually cause fibrosis and cirrhosis, which require liver transplantation^6, 7^. Several factors contribute to the pathogenesis of NAFLD. Unlike the ‘two-hit hypothesis,’ the ‘multiple hit’ concept addresses the interaction of multiple factors contributing to the pathogenesis of NAFLD. These include genetics^8^, environment^9^, lifestyle^10^, insulin resistance^11^, adipocyte differentiation^12^, and the intestinal microbiota^13^. Among the several factors contributing to NAFLD, environmental chemicals play a major role in triggering the pathogenesis of NAFLD^14–16^.

As the most widely produced and prevalent environmental chemical worldwide, bisphenol A (BPA) is a potent endocrine disruptor^17^. Because of its ubiquity in nature and consumer goods, humans and non-human organisms are exposed to this chemical at early developmental stages, leaving behind long-lasting health effects in present and future generations ^18–21^. Several epidemiological studies have found a link between BPA exposure and metabolic disorders, including cardiovascular disease, diabetes, and liver disease^22–24^. There is a positive correlation between BPA levels in urine and obesity, insulin resistance, ALT levels, and hepatic steatosis index levels (HSI)^25–27^. Both *in vitro* and *in vivo* studies showed that BPA increases *de novo* fatty acid synthesis by HepG2 cells and enhances hepatic triglyceride (TG) synthesis in mice^28–30^. BPA promotes excessive reactive oxygen species (ROS) production to advance liver pathogenesis by promoting mitochondrial dysfunction and lipoperoxidation in hepatocytes ^31–33^.

Biologically, sex remains an important variable in the pathogenesis of metabolic disorders^34, 35^. Evidence suggests that direct BPA exposure can promote sex-specific hepatic dysfunction^36, 37^. By establishing the heritable epigenetic memories in germ cells, BPA not only causes adverse health outcomes in the immediate generation exposed to it but also promotes transgenerational health outcomes in unexposed generations^38–42^. We and others have shown that medaka fish can be an animal model for studying human NALFD^43, 44^. Medaka provides additional benefits to understanding the heritability of this disease phenotype across generations. We have reported the heritability of NAFLD up to five generations and sex-specific NAFLD-NASH phenotypes in medaka fish caused by ancestral exposure to BPA^44^. It is unclear whether the transcriptional signatures of medaka NAFLD-NASH phenotypes are similar to human NAFLD-NASH patients. To compare transcriptional signatures associated with NAFLD between human NALFD and BPA-induced NAFLD in medaka fish, we analyzed bulk RNA seq data obtained from the liver of male and female medaka fish that developed sex-specific NAFLD phenotypes and compared them with publicly available human NAFLD and NASH transcriptional signatures. Using a system biology approach, we identified gene sets (biomarkers), transcription factor-driven pathways, and sex-specific up- and downregulated genes and their pathways in fish and human NAFLD-NASH.

## Materials and Methods

### Animal care and ancestral BPA exposure, BPA lineage maintenance, and sample collection

As a relevant animal model for human NAFLD, the present study used medaka fish, which shows liver phenotypes comparable to those of humans^44, 45^. A four-month-old medaka from the BPA and control lineage that developed transgenerational NAFLD-NASH phenotypes was used to determine the transcriptional landscape and NAFLD-NASH genes profile. The maintenance, procedure for exposure, and euthanization were approved by the Institutional Animal Care and Use Committee (IACUC) of the University of North Carolina Greensboro as previously published^44^. In the laboratory, medaka fish were reared in 20 L glass aquariums on a 14 hour:10 hour light cycle, recirculated water with 25% water exchanged every four hours at 26 ± 1 °C, and fed Otohime granular food and brine shrimp (*Artemia nauplii*) twice daily. Every exposure group had three biological replicate tanks, and within each biological replicate group, embryos were collected from three to five separate breeding pairs. To generate fish with transgenerational liver phenotypes, The third generation (F2) was used after initial BPA exposure at the first generation (F0). Bisphenol A exposure can affect metabolic and reproductive health of almost all vertebrates at various concentrations ^46,22, 47–49^. The concentration of BPA (10 μg/L), which is relevant to the environmental levels in many regions of the world, was used as a test concentration^50, 51^. BPA exposure solutions were prepared and measured using mass spectrometry as described elsewhere^52, 53^. The exposure of BPA started after eight hours post-fertilization stage (hpf) and continued until day fifteen after fertilization (dpf). The BPA exposure occurred only at the first generation (F0) for first 15 days of life and never thereafter. Fish were reared in clean water with no further exposure until F2 generation. The exposure window of the first 15 days includes a critical period of sex determination^54^ and liver differentiation^55^ in medaka but excludes the embryonic stem cell differentiation phase in medaka^56^. The uptake of BPA was 20 pg/mg egg/day^44^, which is less than the daily intake of BPA by humans^57^. The measured concentration of BPA was within <10% of the calculated concentration throughout the experiment. The evidence that embryonic BPA exposure (10 μg/L) at F0 generation (ancestors) leads to NAFLD, fertilization defects, and increased embryo mortality in subsequent unexposed generations has been previously demonstrated by our group^44, 58, 59^. At 120 days of age, six pairs of fish from the initial (F0) generation were bred to produce F1 offspring. The same mating methodology was used to produce the next F2 generation (the third generation, first transgenerational). Six males and nine females were used from each lineage (control and BPA). The experimental fish were euthanized with MS-222 (250 mg/L) at the age of four months, and the liver samples were collected from the fish.

### RNA-seq library preparation, RNA sequencing, and data analysis

The liver of control and BPA lineage fish (9 females and 6 males) were used for total RNA extraction by using Quick RNA/DNA Miniprep Plus Kit (#D-7003, Zymo Research, CA, USA) according to the manufacturer’s protocol as previously described^60^. RNA quality was tested by bleach gel electrophoresis^61^, and the quantity was determined by Nanodrop 2000 and Qubit (Thermofisher, Waltham, MA). The RNA of the liver from three fish was pooled to make one biological replicate per group for RNA sequencing. Transcriptome libraries were prepared using NEBNext Ultra II RNA Kit and manufacturer’s protocol. The libraries were sequenced on Illumina Novaseq 6000 (Novogene Corporation, CA, U.S.A.) using a 150 bp paired-end sequencing strategy (short-reads), producing 20–40 million reads per biological replicate. Bioinformatics analysis was performed by using the Longleaf clusters of Supercomputing Facility of the University of North Carolina Chapel Hill. The reads were first preprocessed with Fastp 0.23.2^62^, an ultra-fast all-in-one FASTQ preprocessor, which performs quality control, adapter trimming, quality filtering, per-read quality pruning, and many other operations with a single scan of the FASTQ data. The processed reads were then mapped to the medaka genome (Oryzias_latipes.ASM223467v1) using STAR 2.7.7a^63^. Finally, DESeq2 v1.34.0 was used to do the differential expression analysis^64^. Pathway and network analysis done by using Cytoscape^65^ Shiney GO^66^, and R package^67^.

### Comparative analysis of gene sets: NAFLD/NASH caused by ancestral BPA exposure vs publicly available NAFLD/NASH datasets

A set of DEGs established by direct BPA exposure in mice was obtained from the previously published publicly available data set (NCBI accession number PRJNA529277)^68^. Human patient datasets were collected by compiling data set of GSE89632, GSE99010, GSE52748 and followed by previously published database^69^ and GSE48452^70^. Overlapping DEGs between BPA lineage fish and human patient group were selected and illustrated by using VENNY (http://bioinfogp.cnb.csic.es/tools/venny/index.html). Throughout comparisons, expression data from BPA lineage livers were compared with those from controls, unless specifically mentioned. The transcriptome database has been submitted to NCBI as GSE252744 (token# oruzwusypfezxmp)

## Results

### RNA seq revealed sex-specific transcriptional alterations and potential biomarkers of transgenerational NAFLD induced by ancestral BPA exposure

A bulk RNA sequencing was performed to identify transcriptional alterations in the liver. Global transcriptome screening identified 11,928 and 16,826 differentially expressed transcripts in the liver of BPA lineage males and females compared to the control group. Based on the volcano plot, the livers of the BPA lineage males (Supplementary Figure 1A) displayed a significantly decreased number of unique DEGs than the livers of the BPA lineage females (Supplementary Figure 1B). Using strict selection criteria (5<log2FC< −4 and for females and 2.5 <log2FC< −2 for males), mRNA biomarkers associated with the transgenerational NAFLD phenotype were identified in males (Figure 1A) and females (Figure 1B) of the BPA lineage. As compared to the control group (Figure 1A), *elovl5, igfbp1, tlr5, hck*, and a*pcs* were significantly upregulated, and *elovl1, fbxo4, invs, prkaca,* and *pheta2* were significantly downregulated in the liver of the BPA lineage males. The top 10 up- and down-regulated genes associated with direct exposure to BPA in the male liver were provided in Supplementary Table 1A and B^71^. In the female liver, *vtg3*, *fabp7*, *cacna2d4*, *esr1*, *aldh18a1*, *mttp*, *fas*, and *pycr1* were significantly upregulated while *ppar*α*, acox3*, *cdlk2*, *enpep*, *trak1* and *ugt3a1* were significantly downregulated in the BPA lineage than the control lineage (Figure 1B). A list of significantly up- and down-regulated DEGs in females due to direct BPA exposure was provided in Supplementary Table 1C and D^71^. As the metabolic pathways are common among medaka, mice, and humans^43^, the DEGs (biomarkers) induced by BPA exposure (direct, intragenerational) in the mouse liver were compared between the two vertebrate species-mice and medaka. Results suggest that transgenerational liver disease biomarkers are significantly different from those caused by direct exposure to BPA.

**Figure 1.**
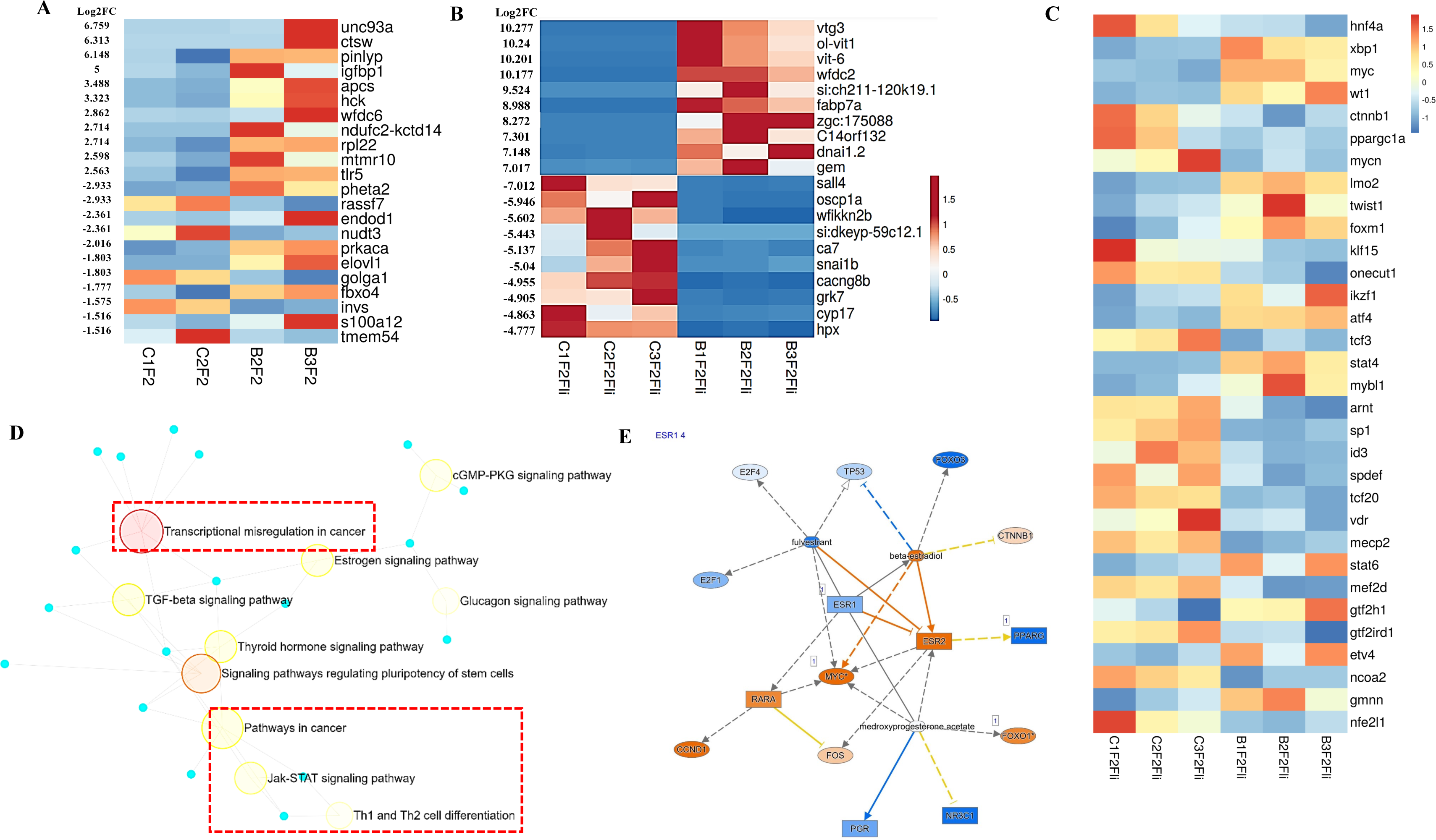
Liver histology of male and female liver of BPA lineage. (A) Control male (B) BPA lineage male and (C) Control female, (D) BPA lineage male. RNA seq determined the top ten up and down regulated genes in (E) male liver and (F) female liver of BPA lineage compared to control. (G) Transcription factors found in the female liver of BPA lineage. (H) Transcription factors drive pathways in the liver of BPA lineage females. (I) Estrogen receptor-mediated gene activation in the liver of BPA lineage female.

### Transcription factors-mediated cancerous pathways were found in the livers from BPA lineage females

To gain insight into how transcription factors play a role in BPA-induced transgenerational NAFLD, we examined the expression of several transcription factors and their roles in triggering disease-specific pathways in the liver of BPA lineage medaka. In the livers of BPA lineage male, *srf, sall1*, *rad21*, *elf1*, *mef2*c were collectively formed transcription factors (TFs) network controlling expression of genes associated with NAFLD phenotype (Supplementary Figure 2A and Supplementary Table 2). TF networks and their expression profiles in the liver of BPA lineage females were shown in Supplementary Figure 2B and heatmaps (Figure 1C), respectively. In the liver of the BPA lineage females, transcription factors, mainly *xbp1, myc, atf4, stat 4, and stat 6* were upregulated, whereas *hnf4a, ppargc1a, klf5, sp1*, and *vdr* were downregulated (Figure 1C). The number of disease-specific TFs found in the livers of BPA lineage females was significantly increased compared to the livers of BPA lineage males. Transcription factors found in the female livers of the BPA lineage were associated with transcriptional dysregulation in cancer, estrogen signaling pathways, Jak-STAT signaling pathways, Th1 and Th2 cell differentiation, and TGF beta signaling pathways in cancer (Figure 1D). Ingenuity pathway analysis (IPA) determined 50 cancerous genes (Supplementary Table 3) expressed in the female livers of the BPA lineage compared to the control lineage. Estrogen receptor-mediated (*esr1* and *esr2*) activation of *myc*, *rara, and ccnd1* and downstream activation of the cross-linking network were provided in Figure 1E and Supplementary Figure 3.

### Both male and female livers of the BPA lineage expressed upregulated genes related to the innate immune pathway, LDL/HDL-mediated lipid transport, and downregulated genes related to lipid digestion

To determine the up and downregulated DEGs and their associated pathways contributing to the NAFLD pathogenesis, significantly altered genes in the liver of BPA lineage were screened by considering significance criteria |log_2_FC| > 0.5, and FDR< 0.1. In total, there were 115 and 1012 significantly upregulated genes and 387 and 702 significantly downregulated genes in the liver of BPA lineage males (Figure 2A) and females (Figure 2B), respectively. Out of all upregulated genes, BPA lineage had 11 common in both sexes, 104 male-specific, and 1001 female-specific DEGs (Figure 2A and Supplementary Table 4A). Out of all downregulated genes, 85 were common in both sexes, 302 were male-specific, and 617 were female-specific (Figure 2B and Supplementary Table 4B). As fatty liver disease phenotypes developed in both males and females from BPA lineages, we explored pathways triggered by up- and down-regulated genes common in both sexes. According to KEGG analysis, commonly upregulated genes were enriched in chemokine and chemokine receptor activation, triggering innate immunity (Figure 2C), suggesting immunogenic pathways being impacted. In addition, down-regulated genes common in males and females indicated impact in lipid digestion, mobilization and transport, lipoprotein metabolism, HDL/LDL-mediated lipid transport, and bile acid synthesis (Figure 2D). Supplementary Tables 5 and 6 demonstrated the enrichment of total pathways associated with DEGs in the BPA lineage males and females. The present results indicated that up- and downregulated genes in both sexes of BPA lineage fish contributed to dysregulation of immunogenic and lipid clearance pathways in the liver.

**Figure 2.**
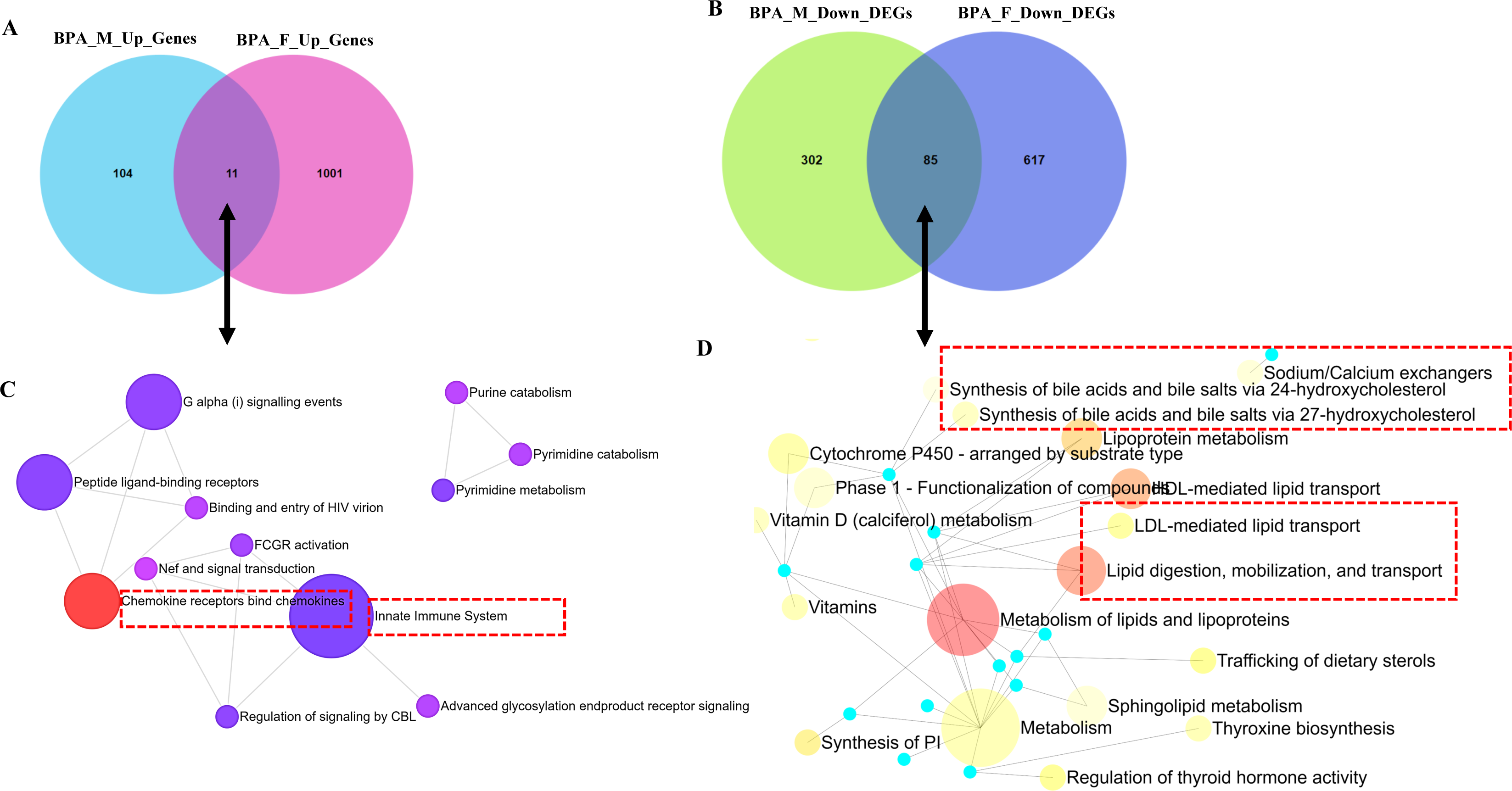
Comparison of upregulated DEGs in the liver of BPA lineage males and females (A) and enriched pathways of upregulated DEGs exhibited (B). Comparison of down-regulated DEGs in males and females (C) and associated pathways (D).

### Sexually dimorphic expression of genes associated with fat metabolism and immunogenic genes in the BPA lineage fish

To ascertain the role of fat metabolizing genes in unequal and sexually dimorphic fat deposition in the liver of the BPA lineage fish, we screened for the expression pattern of DEGs associated with cholesterol metabolism, fatty acid transport, lipogenesis, and lipolysis. Heatmap was used to demonstrate the expression pattern of fat metabolizing genes in the liver of BPA lineage males (Figure 3A) and females (Figure 3B). The expression of genes encoding enzymes for the lipogenic cycle, mainly *sqle*, *mttp*, *cers5,* and *aco1* was significantly downregulated in the liver of the BPA lineage males compared to the control females (Figure 3A). However, lipogenic genes such as *scd, mttp, and abcg1* were significantly upregulated in the liver of BPA lineage females compared to controls, except for *pnpla3* (Figure 3B). This suggested that lipogenesis was elevated in females’ livers than males of the BPA lineage. In contrast, genes encoding enzymes controlling the lipolytic cycle were significantly downregulated in both males and females of the BPA lineage. Downregulated lipolytic genes included *acadm*, *cpt1ab*, and *ppar*δ in the males and *ppar*α, *ppar*δ, *cyp 450*, and *crot* in the females, suggesting that the lipolytic cycle was disrupted in the liver of BPA lineage fish. The mRNAs for *srebf1*, *srebf2*, *ppar*γ, and *pparggc1a* were significantly decreased in the liver of BPA lineage females compared to the control females. Genes associated with fatty acid transport mechanisms, such as *osbp, cetp, lrpap1,* and *slc27a1* in the male liver and *apoab* and *fabp3* in the BPA lineage female livers, were significantly downregulated, suggesting restricted fat release from the liver in the BPA lineage fish. In contrast to aberrant expression of fat transport genes, the receptor protein CD36, which controls the intake of extrahepatic fat granules, was significantly increased in the liver of BPA lineage females compared to control females. The receptor protein *cd36*, which controls the intake of extrahepatic fat granules, was significantly upregulated in the liver of BPA lineage females, suggesting an increased intake of fatty acids rather than an effective fat clearance process. In the liver of both BPA lineage males and females, the DEGs associated with cholesterol metabolism were downregulated, but *soat2* was upregulated in the liver of the BPA lineage males compared to control males. BPA lineage males showed downregulation of *acot7, elovl6*, and elovl5, while elovl1 was upregulated. The expression of *tlr5, tnfaip3, cxcl12, hla-dqa1*, *rsl24d1,* and *cxcr4* were considerably increased in the liver of BPA lineage males compared to control males (Figure 3A). *cxcr4, cxcl12* were upregulated and *irf3, caspase3, caspase8, ap1m2, il10*, *tlr8* and *irak1bp1* were upregulated in the liver of BPA lineage females compared to controls (Figure 3B).

**Figure 3.**
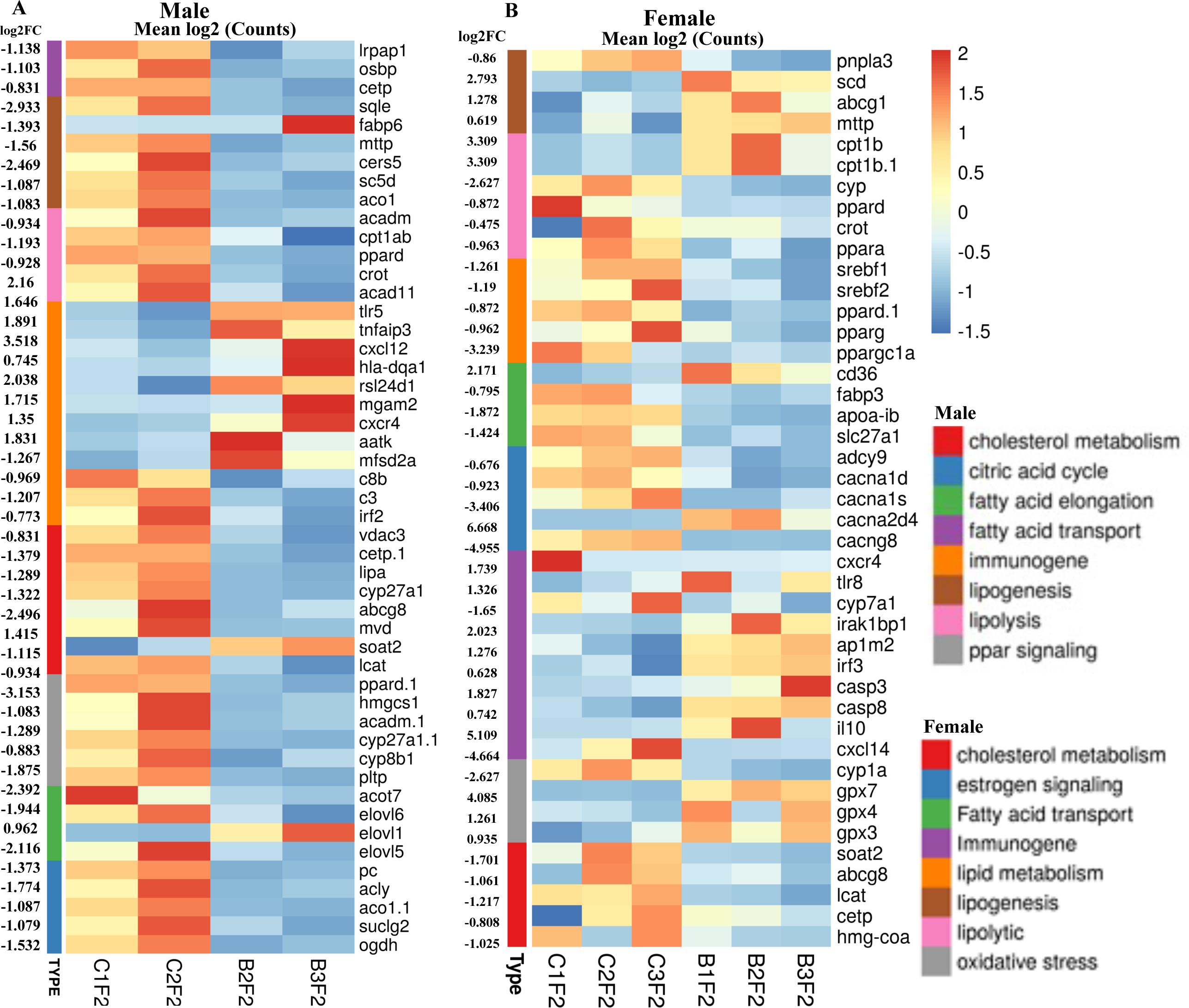
Heat maps showing fat metabolism genes, including cholesterol metabolism, lipolysis, lipogenesis, and fatty acid transport, were displayed with sex-specific expression patterns. Differentially expressed genes in the male liver (A) and female liver (B) are associated with fat metabolism are shown. Color codes indicate treatments and significantly affected pathways.

### The liver of BPA lineage females showed abnormal expression of genes associated with estrogen signaling, epigenetic processes, aberrant cell cycle regulation, activation of kinases, and cytokines

In the BPA lineage females, estrogen signaling pathway genes such as *adcy9, cacna1s, cacna1d,* and *cacng8* were downregulated except for *cacna2d4* (Figure 3B). In addition, genes encoding for estrogen-dependent receptors, including *esrp1, esr1, cav2,* and *cavin1b*, were significantly upregulated, whereas esr2 and *esrp2* were downregulated (Figure 4A). A list of genes involved in estrogen receptor-mediated activation of downstream genes is shown in Supplementary Figure 3. To explore the involvement of epigenetic processes in disease phenotype, we examined genes encoding enzymes involved in epigenetic modification in the livers of the BPA lineage females, which showed increased NAFLD phenotype. The genes related to DNA demethylation, mainly *tet1*, *tet2*, and *tet3*, were significantly downregulated, and all subtypes of *dnmts*, *pcna*, and *uhrf1* were significantly upregulated in BPA lineage females than control females (Figure 4B). The genes encoding histone deacetylation, mainly *hdac1, hdac5, hdac7b, hdac11, hdac4* were significantly downregulated, but *hdac9b, hdac8, hdac3, hdac12, hdac6,* and *sirt* were upregulated (Figure 4B). This suggested that epigenetic dysregulation in the liver of BPA lineage females due to ancestral BPA exposure. The genes encoding kinases, including *map3k21, ripk2, mapk1, and mtor* were differentially expressed (Figure 4C), and cytokine genes were significantly upregulated in the liver of BPA lineage females than control livers (Figure 4D). The cell cycle regulation genes *wee1*, *aruka*, *rrm2*, and *rrm2* were significantly upregulated in the liver of BPA-lineage females compared to control females. The cell cycle regulation genes *wee1, aruka, rrm2*, *yme1|1* and *rrm2* were significantly upregulated in BPA-lineage female livers compared to control female livers (Figure 4E) and these genes were associated with G1/S, G2/M transition, APC/C mediated degradation of cell cycle protein, DNA replication (Supplementary Figure 4A).

**Figure 4.**
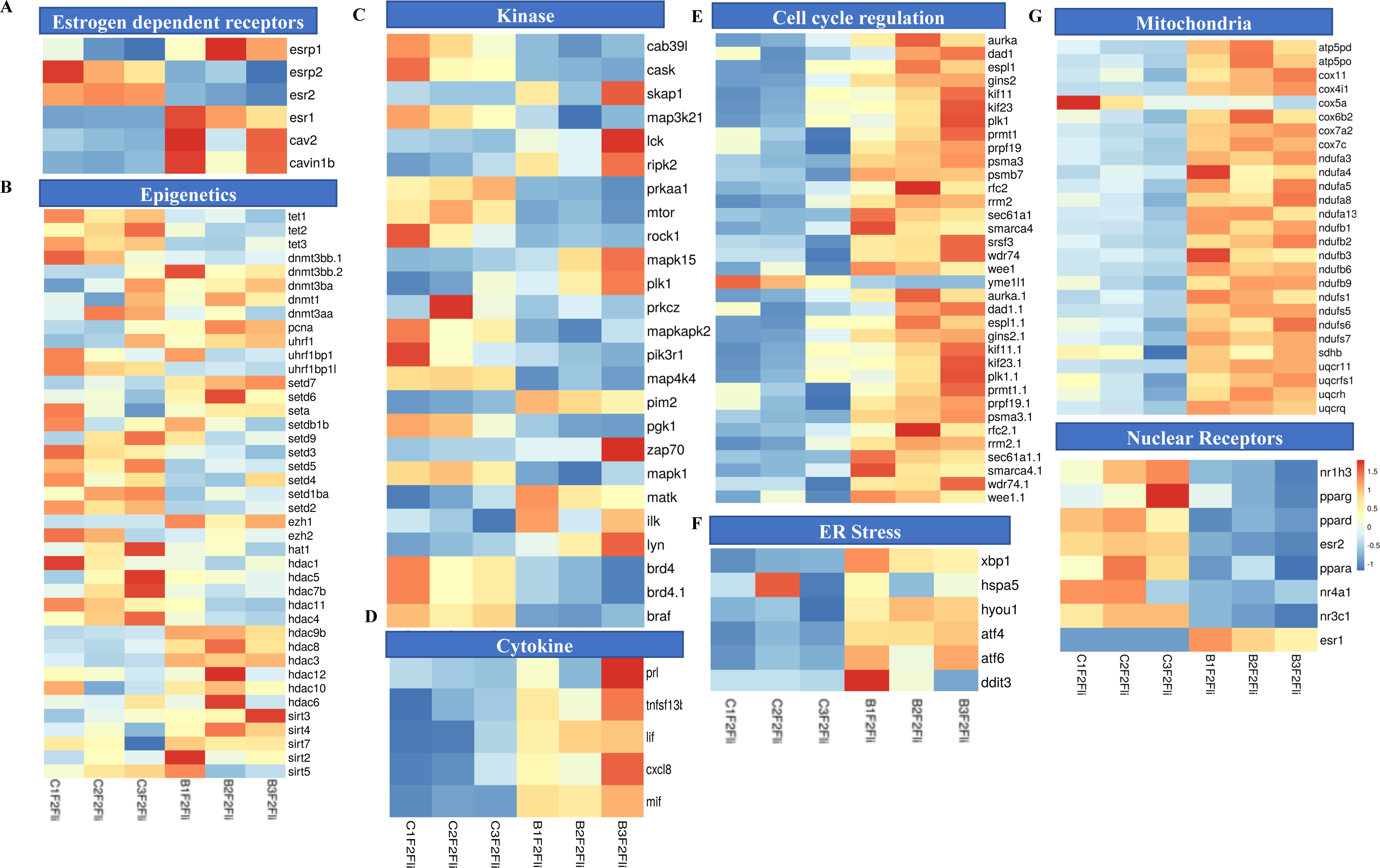
Females in BPA lineage-specific aberrant gene expression in (A) estrogen signaling, (B) epigenes, (C) kinase, (D) cytokine, (E) cell cycle regulation, (F) ER stress, (G) mitochondria, and (H) nuclear receptor compared to control.

### Impact on ER and mitochondrial genes in developing oxidative stress response in the liver of BPA lineage females

When compared between females in the BPA and control lineages, endoplasmic reticulum-mediated stress response genes, mainly *Xbp1*, *hspa5*, *hyou1*, *atf4, atf6,* and *ddit3* were significantly upregulated in the liver of the BPA lineage females (Figure 4F). Among all ER stress genes, *atf4, hspa5, ddit3,* and *hyou1* were upstream hub genes that control many downstream genes, such as *mapk14, jun, ankrd1, and fgf19* (Supplementary Figure 4B). These genes are associated with the production of ROS, unfolded protein, and ER-mediated stress (Supplementary Figure 4B). To investigate mitochondrial gene expression associated with energy metabolism, we found a total of 25 DEGs linked to the electron transport chain (ETC). Among the genes encoding protein complex in ETC, *cox6b2, cox7a2, nufa-3,-4,-5,-8, nduf-1,-2,-3,-6,-9* were significantly upregulated, while *cox5a* was only downregulated in BPA lineage females, suggesting dysregulation of mitochondrial energy metabolism pathway ( Figure 4G and Supplementary Figure 5 and 6). The genes encoding nuclear receptors were significantly downregulated except *esr1* (Figure 4G). Furthermore, oxidative stress-induced genes, mainly *gpx7, gpx4,* and *gpx3* were highly upregulated in BPA lineage females (Figure 3B). The present results indicated activation of oxidative stress in the liver of BPA lineage females, possibly mediated by mitochondrial and endoplasmic reticulum-mediated stress.

### DEG-mediated pathways, Gene Ontology (GO), and IPA analysis revealed disease-related pathways triggered in females from the BPA lineage

In the BPA lineage males, upregulated genes in the DEG list were enriched in Th1 and Th2 cell differentiation, cysteine and methionine metabolism, the intestinal immune network for IgA production, and thiamine metabolism pathways (Figure 5A), suggesting dysregulation of immunogenic response. As revealed by the KEGG pathway analysis, male DEGs were associated with metabolic pathways, protein processing in the endoplasmic reticulum, cholesterol metabolism, and PPAR signaling (Supplementary Figure 7A). In the BPA lineage females, upregulated DEGs were typically associated with cell cycle and checkpoint, p53-independent DNA damage, unfolded protein response by activation of *XBP1*, respiratory electron transport, and apoptosis regulation pathways, suggesting cellular stress resulting in the development of severe phenotypic traits in females from the BPA lineage (Figure 5B and Supplementary Figure 7B).

**Figure 5.**
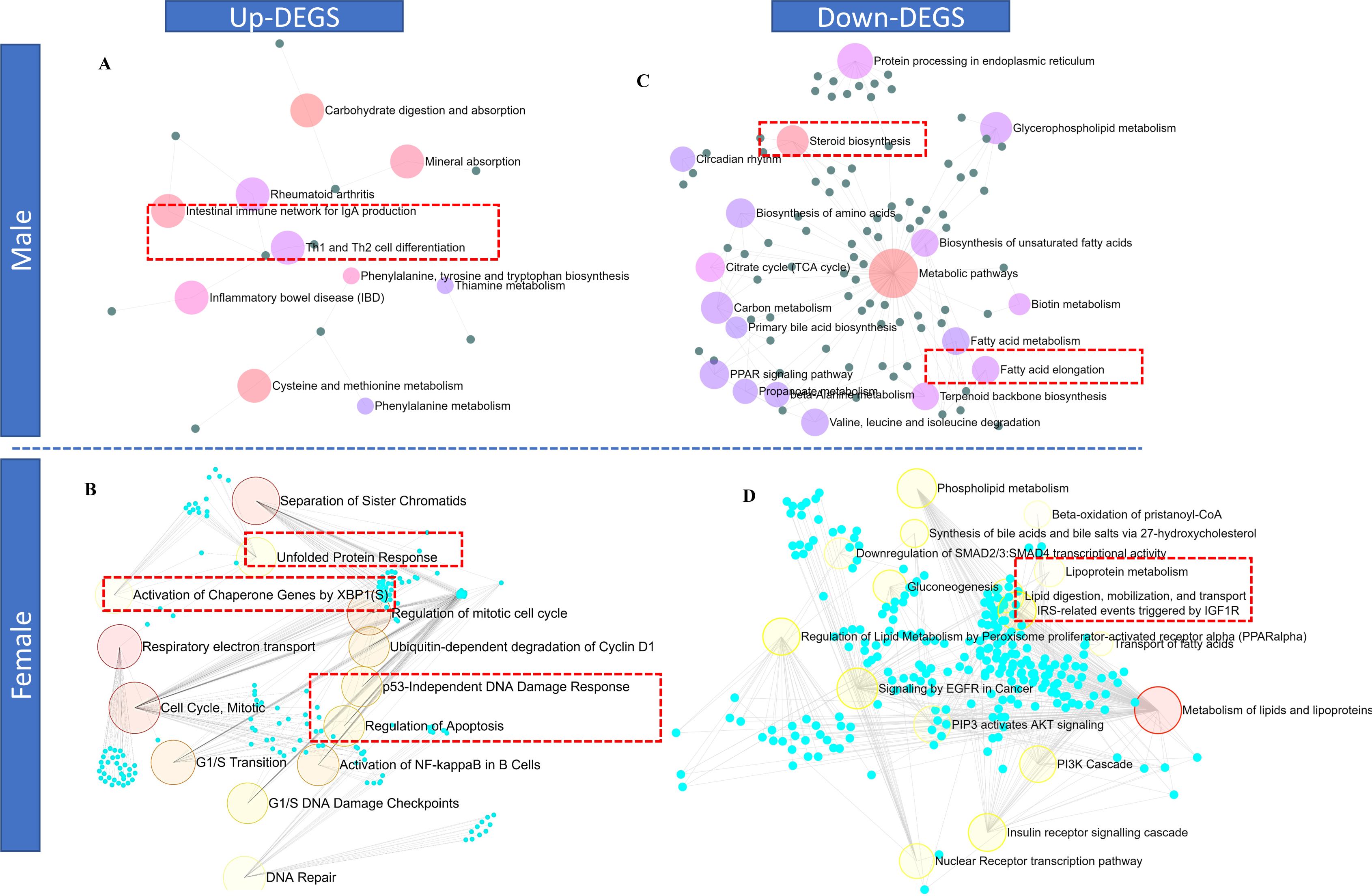
The upregulated genes in the liver of BPA lineage male (A) showed activation of immunogenic pathways while the liver of BPA lineage female (B) showed enrichment of unfolded protein, P53 DNA damage response, and activation of chaperon genes. The downregulated genes in the liver of BPA lineage male (C) showed enrichment in steroid biosynthesis, fatty acid elongation but lipoprotein metabolism, lipid digestion, phospholipid metabolic pathways were enriched in the liver of BPA lineage female (D).

Downregulated DEGs in the males from the BPA lineages were mainly linked to fatty acid metabolism, steroid biosynthesis, fatty acid elongation, and metabolic pathways (Figure 5C). In contrast, beta-oxidation, *PPAR*α regulation, lipid and lipoprotein metabolism, gluconeogenesis, and transport of fatty acid pathways were associated with downregulated DEGs, indicating perturbed lipolysis and fatty acid clearance mechanism in the liver of BPA lineage females (Figure 5D). In total, 15 pathways common to both males and females, 2 pathways specific to males, and 198 pathways specific to females were found to be activated in the liver of BPA lineage fish, suggesting sex-specific associations of disease pathways in liver disease development (Supplementary Figure 7C). Metabolic pathways, cholesterol metabolism, fatty acid elongation, and PPAR signaling pathways were found to be mutually triggered due to ancestral BPA exposure regardless of sex (Supplementary Figure 7C). According to KEGG pathway analysis, NAFLD, cytokine-cytokine receptor interaction, leukocyte migration, antigen processing and presentation, Jak-STAT signaling and MAPK, AMPK, Insulin, thyroid hormone signaling pathways were found to be associated with up- and downregulated DEGs, respectively in the liver of BPA lineage females compared to control females (Supplementary Figure 8).

Next, we performed gene ontology analysis with top-ranked significant DEGs in males and females from the BPA lineage. In the liver, Gene Ontology Enrichment (GEO) analysis revealed a total of 12 (male) and 94 (female) molecular functions (MF), 85 (male) and 678 (female) biological processes (BP), and 23 (male) and 183 (female) cellular components (CC) in BPA lineage fish (Supplementary Figure 9 and 10). In the liver of BPA lineage males, cholesterol metabolic and biosynthetic process (GO:0016126 and GO:0008203) fatty acid synthase and elongase activity (GO:0004312 and GO:0009922) were significantly enriched. Cellular protein modification and metabolism (GO:0006464 and GO:0044267), ribosome biogenesis (GO:0042254), kinase activity (GO:0016301), mitochondrial membrane (GO:0005743 and GO:0031966), focal adhesion (GO:0005925) were detected in the liver of BPA lineage females. The results of IPA pointed out disease-specific canonical pathways, such as liver hyperplasia, hepatocellular carcinoma, liver steatosis, and liver necrosis in the liver of BPA lineage females (Supplementary Figure 11). Adipogenesis, triacylglycerol degradation, and retinol biosynthesis were found in the liver of BPA lineage males (Supplementary Figure 12), whereas liver necrosis, and hepatocellular carcinoma were found in the liver of BPA lineage females as per IPA. Upstream regulators *hnf4a, tp53,* and *xbp1* expressed in the liver of the BPA lineage females (Supplementary Figure 12).

### Comparative analyses of biological processes altered in the liver due to direct vs indirect BPA exposure across humans, mice, and medaka

To investigate liver-specific pathways associated with direct and indirect BPA exposure, a comparison of KEGG pathway analyses was performed, as shown in the Venn diagram. The results showed that directly exposed mice and indirectly exposed medaka shared eight common pathways induced by BPA (Figure 6A). In males, metabolic pathways, cholesterol metabolism, PPAR signaling pathway, cholesterol metabolism, lysosome, and carbon metabolism pathways were common in databases obtained from direct and indirect BPA exposure. In females, KEGG pathway analysis (Figure 6B) found directly exposed mice and indirectly exposed medaka shared ninety-two common pathways induced by BPA. Common pathways in females represented oxidative phosphorylation, metabolic pathways, ribosomes, chemical carcinogenesis, and proteasomes in the liver. However, MTOR, AMPK signaling, insulin resistance, nonalcoholic fatty liver disease (NAFLD), and pathways in cancer were uniquely triggered in the female liver due to ancestral BPA exposure. Results indicated that in females, similar pathways were affected by the direct and indirect exposure to BPA. Compared to BPA lineage males and females, human and mouse transcription factors associated with steatosis were found most commonly in the liver of BPA lineage females (Supplementary Table 7), suggesting disease-specific transcription factors in humans and mice were significantly expressed in the female liver of BPA lineage.

**Figure 6.**
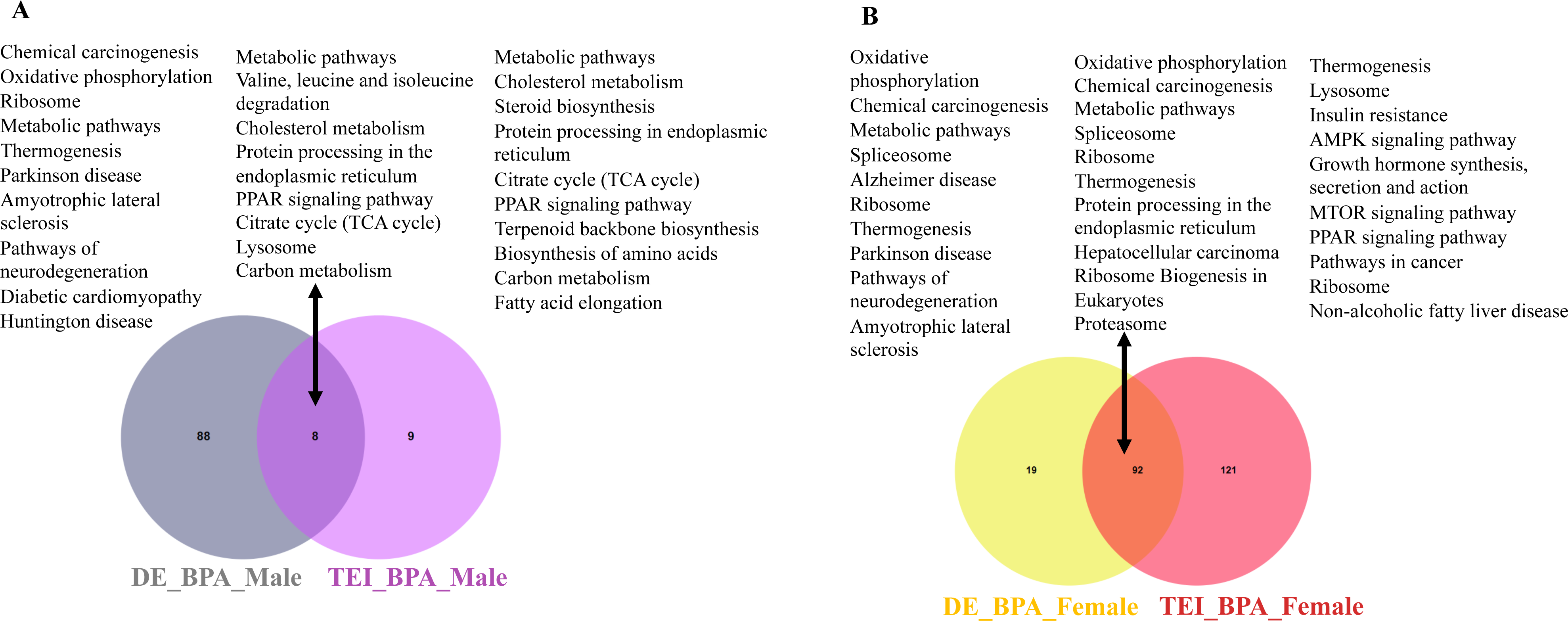
Comparative analysis of pathways triggered due to direct exposure (mice) and indirect exposure (medaka) of BPA exposure in males (A) and females (B) across the species.

### Comparison of human NAFLD-NASH signature genes with those in female medaka from BPA lineage

Ancestral exposure led to NAFLD-NASH phenotypes in the liver of BPA lineage females^44^. Here we screened for NAFLD-NASH signature genes in the livers of BPA lineage females and compared them with gene signatures associated with NAFLD-NASH in human patients. The BPA lineage females and human NAFLD patients shared 88 DEGs (Figure 7A). Additionally, 20% of DEGs were mutually upregulated between medaka and human NAFLD patients (Figure 7B). Among the total shared DEGs, 27.69% of DEGs were mutually downregulated. The total mutual DEGs triggered in the female liver of BPA lineage and human NAFLD patients were illustrated in Figure 7C. A total of 52.31% of DEGs expressed in BPA lineage females were expressed in human patients (Figure 7B). The commonly expressed DEGs in medaka and human NAFLD patients were found to be associated with immune response, cholesterol and lipid metabolic process (Figure 7D). The common downregulated genes were *abcb11, aldh6a1, igf1, cyp1a1, and apof*. Among the shared upregulated genes, *atf3, cxcr4, plin1, gins1, and alpk2* were common in BPA-lineage females and human patients (Supplementary Table 8). In BPA lineage females, gene signaling networks identified *hsp90ab1*, *myc*, *ctnb1*, *tp53*, and *mitf* as potential upstream genes involved in human NAFLD (Supplementary Figure 13). In total, 14 pathways associated with NAFLD pathogenesis were found to be common between BPA lineage females and human NAFLD patients (Figure 7E). Among the common pathways, PPAR signaling and metabolic pathways were highly significant in human NAFLD patients (Figure 7F) and BPA lineage females (Figure 7G), respectively. Genes associated with gluconeogenesis, lipid metabolism, mitochondria, ceramide metabolism, insulin pathway, inflammatory response, cell adhesion, coagulation, cytokines controlling liver fat, and advanced liver disease in humans were comparable to liver genes of BPA lineage females suggesting an association of similar DEGs in NAFLD pathogenesis (Supplementary Table 9). In addition, 39 common DEGs were detected in patients with NASH and BPA lineage females, and only 15 DEGs showed ancestral BPA-specific expression (Figure 7H). Among common NASH-specific DEGs, *igf1* was found to be an upstream regulator connected to downstream genes, including *hnf1a, cyp7a1, mmp13*, *fbn1*, and linked to adipogenesis in the liver (Supplementary Figure 14). The synthesis of extremely long-chain fatty acyl CoAs, fatty acid triacylglycerol and ketone body metabolism, bile acid, and salt metabolism were found to be linked to the common NASH genes network (Supplementary Figure 15).

**Figure 7.**
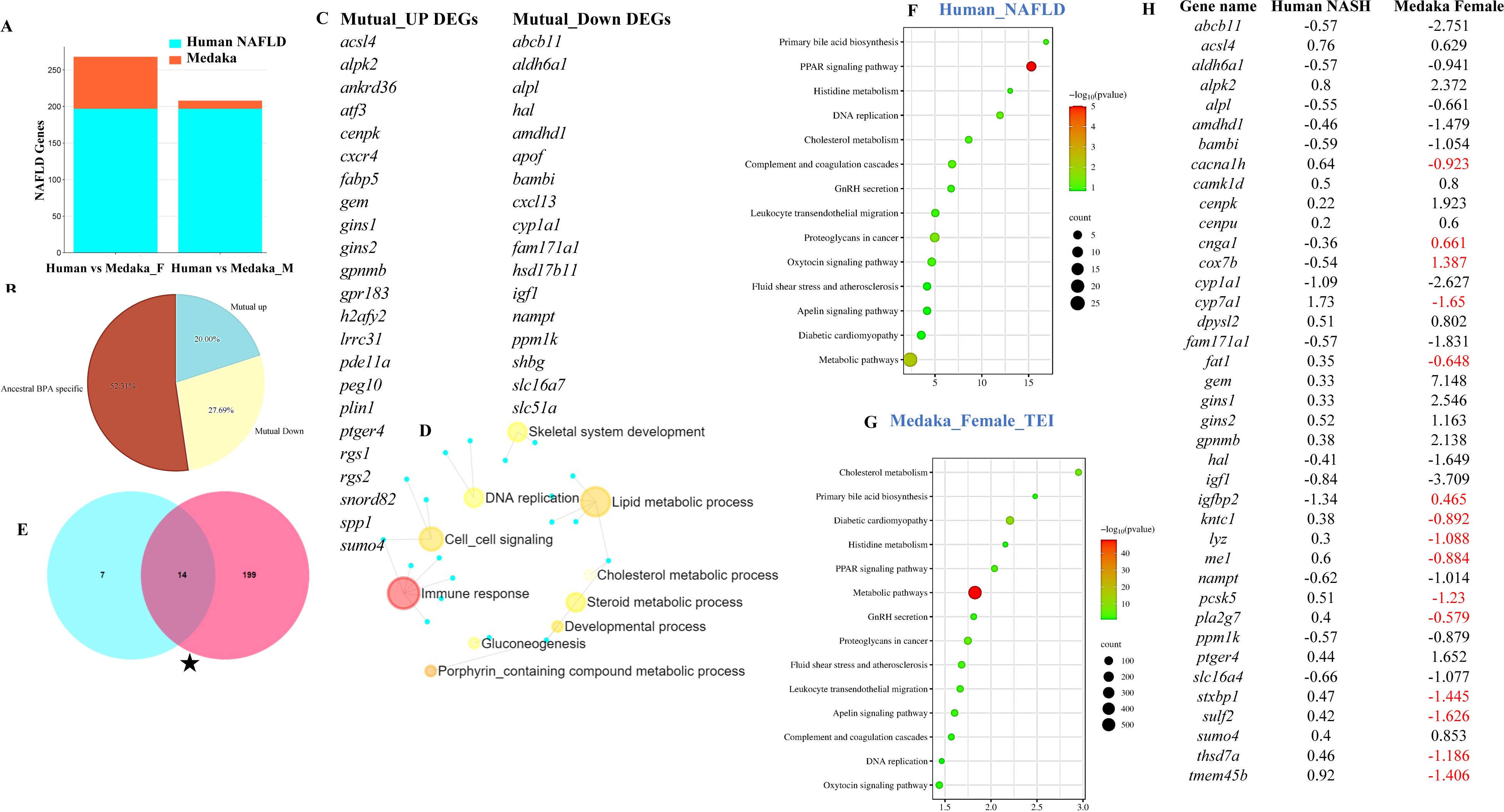
Comparative analysis of DEGS found in human NAFLD and NASH patients and BPA lineage liver of males and females. (A) Barplot showing common DEGs in human NAFLD genes and the liver of BPA lineage female and male. (B) Percentage of mutual up and down-regulated ancestral BPA exposure-specific DEGs in the female liver. (C) Gene signaling network of common DEGs in human NAFLD patients and females’ liver of BPA lineage. (D) Comparison of pathways in human NAFLD patients and BPA lineage females. (E) Total common pathway enrichment in human NAFLD (E) and medaka female (F). (G) Total common NASH-specific genes in human patients and BPA lineage females. (H) Gene signaling network of common NASH genes in BPA lineages.

## Discussion

BPA can alter metabolic health in humans and non-human organisms; however, excluding BPA from consumer goods may not indicate environmental safety. BPA exposure may still cause transgenerational liver disease in descendants who will not experience BPA exposure in the future. This study demonstrates ancestral BPA-specific transcriptional signatures linked to transgenerational NAFLD in the third generation (F2) induced by ancestral embryonic BPA exposure at the first generation (F0). We found that ancestral BPA exposure led to the development of the NAFLD phenotype and the activation of different disease pathways in its progression to NASH by activating genes found in human NAFLD-NASH patients. Interestingly, the pathways linked with BPA exposure-induced liver disease caused by direct BPA exposure differed from those associated with ancestral BPA exposure, suggesting the involvement of novel mechanisms in developing liver diseases in unexposed descendants of exposed ancestors.

Global transcriptomic alterations were determined in the liver of both males and females, but female livers showed massive transcriptional alterations associated with DEGs and disease pathways. Several biomarkers, mainly *bend7, aldh18a1, pycr2,* and *cacna2d4* identified in the liver of BPA-lineage females, have been described as “implicated in the activation of cancerous pathways” ^72–76^. Interestingly, biomarkers linked with direct and ancestral BPA exposure were substantially different, indicating the involvement of novel pathways in the pathogenesis of BPA-induced transgenerational NAFLD.

Transcription factors, primarily *sp1*, *elf1*, *klf5*, and *srebf2* were found in male livers, whereas upregulated *myc, xbp1, atf4, stat6*, and downregulated *hnf4a, ppargc1a*, and *ctnnb1* were found in female livers of BPA lineage. Elevated levels of *xbp1* and *atf4* are linked to inflammation and fibrosis in humans and mice that are undergoing NAFLD^77, 78^. Thus, overexpression of *xbp1* in the liver of BPA lineage females could have triggered lipogenesis and a UPR-driven fibrogenic cascade ^79, 80^. Mutant mice for *atf4* exhibited recovery in liver steatosis phenotype by AMPK-dependent inhibition of fatty acid synthase^81^. Elevated levels of *ATF4* found in NASH patients and depletion of *ATF4* displayed a protective role in advanced liver phenotype^81^. A previous study found that the activation of *myc,* one of the most commonly activated oncogenes, is a critical driver of liver carcinogenesis both *in vitro* and *in vivo*^82^. Yang *et al.* showed an activation of *hnf4a,* resulting in nuclear transcription factor-mediated attenuation of liver fibrosis ^83^. As a result of ancestral BPA exposure, the downregulation of *hnf4a* and overexpression of *xbp1* and *atf4* could have induced liver fibrosis and progressed the NAFLD to NASH phenotype in medaka. It is worth mentioning that BPA induced equivalent expression of *hnf4, xbp1,* and *atf4* in the livers of both immediately ^84–86^ and ancestrally exposed generations. The role of transcription factors activated in the female liver was associated with transcriptional misregulation in cancer, estrogen signaling route, Jak-STAT signaling, and TGF beta signaling system, according to enrichment analysis, and most of the pathways triggered were associated with the advanced stage of liver disease^87, 88^.

In addition to a few common DEGs, we found more sex-specific DEGs between males and females of BPA lineage than in control. DEGs that were identified to be commonly elevated in both males and females were connected to chemokine and innate immune activation. Interestingly, the same BPA lineage (F2 generation) exhibited transgenerational gut dysbiosis^89^, indicating a possible relationship between gut dysbiosis and activation of immunogenic response in the liver of BPA lineage fish, which could have enhanced the inflammation. Mutually expressed downregulated genes in both males and females of the BPA lineage were enriched in LDL and HDL-mediated lipid transport, lipid digestion, and synthesis of bile acids. Studies showed that low-density apolipoprotein receptor (*LDLR*) and high-density apolipoprotein receptor (*HDLR*) play an essential role in the primary route of circulatory lipid clearance in liver cells^90, 91^. Choi *et al.* demonstrated that knockout (*Ldlr^−/−^*) mice were impaired in apoB and apoE-containing lipoprotein clearance and induced steatosis and inflammation in liver^92, 93^. Together with published literature, downregulated genes associated with lipoprotein metabolism could have perturbed the release of fat droplets from the liver in the BPA lineage fish and promoted NAFLD. However, how the heritable epigenetic signature in sperm epigenome controls the differential expression of liver genes and how it is linked to pathogenesis need to be determined in future studies.

As fat-metabolizing genes play a primary role in lipid metabolism of the liver, we screened genes associated with cholesterol metabolism, fatty acid transport, lipogenic, and lipolytic genes in both male and female livers of BPA lineage. We found that most lipogenic genes were downregulated in males, whereas lipogenic genes such as *scd*, *mttp*, and *abcg1* were dramatically elevated in female BPA livers, but *pnpla3* was downregulated. According to the literature, direct BPA exposure increased adipogenesis in the liver by increasing expression of *SCD* and *ABCG1* in vivo^84, 94^, and *ABCG1* was linked to steatohepatitis^95^. *SCD1* gene knock-out mouse study showed a decrease in the lipogenic cycle and fat accumulation in the liver^96, 97^. This suggested that activation of lipogenic genes in the female liver could contribute to abnormal fat accumulation. In contrast, genes linked with the lipolytic cycle in the liver, *acadm*, *cpt1ab*, and *ppar*δ were downregulated in males, whereas *cyp450*, *ppar*δ, *ppar*α, and *crot* were downregulated in females of the BPA lineage. Interesting findings included the up- and downregulation of *cpt* in the male and female BPA lineages compared to controls, respectively. Literature suggested that downregulation of *ppar*α displayed severe hepatosteatosis^98^, and depletion of *ppar*δ exhibited increased hepatic expression of inflammatory cytokines *TNF*α and *IL1*β and fibrosis in response ^99^. Additionally, genes that control beta-oxidation and lipid metabolism, including *cpt1b, srebf1*, *srebf2*, *ppar*γ, and *pparggc1a* were downregulated in liver of BPA lineage females. Previous studies showed that the upregulation of *CPT* genes determines the activation of beta-oxidation pathways ^100^ and direct exposure to BPA decreased *CPT* expression in a previous study^101^. Thus, an increased expression of *cpt1b* gene in the liver of BPA-exposed females was unique, indicating an enhanced mitochondrial lipid catabolism pathway. However, upregulation of *cpt1b* in the liver of BPA lineage females could have triggered the apoptotic mechanism by elevating CD4^+^ T^102^. Ke *et al*. showed that direct exposure to BPA decreased the DNA methylation levels of *Srebf1* and *Srebf2* that increased with lipid synthesis^103^. An ancestral BPA exposure led to downregulated *srebf1* and *srebf2* in the liver of BPA lineage females which might suppress sterol biosynthesis^104^. Instead of *de novo* fat synthesis in the liver of BPA lineage females, upregulation of gene encoding fat translocase protein in hepatocytes, *cd36,* could have induced the uptake of extrahepatic fat through the portal and central circulation^105–107^. Decreased cholesterol biosynthesis in the liver of the BPA lineage fish could have impaired steroidogenic hormonal pathways, leading to reproductive disorders found in the BPA lineage. As a result, dysregulation of fat metabolism could have contributed to abnormal fat accumulation and reduced sterol biosynthesis, leading to reproductive impairment in BPA lineage fish. Future research must explore the epigenetic control mechanisms involved in the transgenerational dysregulation of fat-metabolizing genes in the liver.

Carbohydrate digestion, Th1 and Th2 cell differentiation, cysteine and methionine metabolism were enriched in the liver of BPA lineage males. Pacana *et al*. showed that multiple alterations in methionine metabolism trigger NAFLD pathogenesis^108^. Increased hepatic production of Th1-related cytokines IFN, IL-12, and TNF triggered hepatitis in choline-deficient-diet-fed steatotic mice^109^, as well as elevation of genes toward a Th1 cell differentiation when comparing NASH patients to those without NASH^110^. Transgenerational gut dysbiosis in the same male/ female fish from BPA lineage could have activated Th2 cell activation that could interfere with IL-4/STAT6 axis on insulin sensitivity, leading to the inhibition of *PPARa,* lipolytic gene^111^. Aberrant methionine metabolism and disruption in the lipolytic cycle in the BPA lineage males could have contributed to enhancing fatty acid elongation process and sterol biosynthesis^112^. In the female liver, pathways to P53-dependent DNA damage response, activation of NF-kappa B cells, activation of chaperone genes by XBP1, and dysregulation in the cell cycle were mainly enriched with upregulated DEGs in the BPA lineage females. Several studies showed that direct exposure to BPA induces DNA damage^111^, promotes cell proliferation in c-Myc-dependent manner in estrogen receptor (ER)-α-negative mammary cells^113^, increases the generation of reactive oxygen species (ROS) in the liver ^114^, and induces the formation of DNA adducts^115^. Additionally, abnormal fat accumulation in BPA lineage females could enhance oxidative stress response and apoptotic mechanism by activating caspase and Gpx^116, 117^.

Importantly, the signaling pathway activated by ancestral BPA exposure in the female liver, including disruption of ER homeostasis and the unfolded protein response (UPR), has been linked to lipid biosynthesis, insulin action, inflammation, and apoptosis^118–120^. Oxidative phosphorylation, chemical carcinogenesis, and metabolic pathways were mutually triggered in direct and indirect BPA exposure but in the liver of BPA lineage females. However, pathway enrichment analysis uncovered unique pathways such as AMPK signaling activation (43 genes, FDR −2.92E-06), MTOR signaling (48 genes, FDR −4.56E-05), Rap1 signaling (61 genes, FDR-4.04E-05), pathways in cancer (125 genes, FDR-0.000102067) only active in the female liver of the BPA lineage that has been associated to cancer^121–124^. Taken together, ancestral BPA exposure could have triggered harmful pathways, resulting in a severe liver phenotype.

We found 88 common DEGs associated with NAFLD in BPA lineage females and human NAFLD patients. Among the shared downregulated genes, a study showed that significantly decreased *Cyp1a1* expression increased cholesterol synthesis and reduced intrahepatic lipid accumulation^126^. Other studies found that *abcb11* knockout mice exhibited impaired mitochondrial fatty acid β-oxidation that might exacerbate cholestatic liver damage^127^. The decreased expression of *igf1* was associated with histologic severity of NAFLD^128^. Overexpression of *atf3-*induced hepatic stellate cell activation was found in both *in vivo* and *in vitro* studies, and knockdown of *atf3 alleviated* advanced liver disease^129^. Compared with wild-type mice, mice lacking or overexpressing hepatic ATF3 exhibit decreased or increased RIPK3 expression in severe hepatic steatosis and necroptosis after partial hepatectomy. Based on live cell imaging experiments, *atf3* induces necroptosis instead of apoptosis in cultured hepatocytes^130^. Upregulation of *fabp5* was associated with the activation of hepatocellular carcinoma^131^. In the advanced stage of liver disease, *cxcr4* expression was found to be increased^132^ and is associated with hepatocellular carcinoma^133^. Overexpression of *plin1* increases lipid metabolism and also enhances triglyceride levels^134^. In total, we found 39 genes common between NASH patients and livers of BPA lineage females, and 15 DEGs showed unique ancestral BPA exposure-specific expression patterns. Among common genes, *cnga1, cox7b, igfbp2* were upregulated and *cacna1h, cyp7a1, fat1,kntc1,lyz,me1,pcsk5,* and *pla2g7* were downregulated showing ancestral BPA specific expression patterns. However, the downregulation of *igfbp2* was associated with fatty liver^135^, the upregulation of *cox7b* is linked to the disruption of mitochondrial electron transport change^136^, and the downregulation of *cyp7a1* was associated with the synthesis of cholesterol^137^. Among NASH-associated DEGs, *igf1* was highly downregulated in the liver of BPA lineage female fish compared to human patients. Literature suggested that impairment of IGF1 synthesis results in a worsening state of insulin resistance^138^, and reduced *IGF1* expression caused fibrosis^139^, suggesting *igf1* could have enhanced NASH pathogenesis in the liver of BPA lineage females, a master regulator.

## Conclusion

This study aimed to examine the global changes in transcriptional networks and their connection to the pathogenesis of NALFD and NASH in the grandchildren generation affected by ancestors’ exposure to BPA. In the livers of the BPA lineage females, abnormal cytokine and kinase gene activation that might set off multiple disease pathways were found. Our results indicated that ancestral BPA exposure leads to the activation of several DEGs associated with human NAFLD-NASH patients with alteration of LDL/HDL-mediated lipid transport, metabolism of lipoprotein and lipids, p53-dependent DNA damage response, AMPK, mTOR, and cancerous pathways. Comprehensive transcriptional network analysis identified probable mechanisms of fat accumulation, oxidative stress response mediated by mitochondria and ER, and liver fibrosis in female livers caused by the impact of ancestral BPA exposure.

## Acknowledgments

The authors thank undergraduate researchers for their help in experimental procedures and the Department of Biology, University of North Carolina Greensboro (UNCG) and UNCG Graduate School for their research support to SC.

## Author contributions

Conceptualization: RKB, SC; Methodology: SA, SC; Validation: SA; Formal analysis: SC, SA; Investigation: SC; Writing original draft: SC; Writing-review & editing: SC, RKB; Supervision: RKB; Project administration: RKB; Funding acquisition: RKB.

## References

(1) Machado, M. V.; Cortez-Pinto, H. Non-alcoholic fatty liver disease: what the clinician needs to know. World J Gastroenterol 2014, 20 (36), 12956–12980. DOI: 10.3748/wjg.v20.i36.12956

(2) Liu, Q.; Bengmark, S.; Qu, S. The role of hepatic fat accumulation in pathogenesis of non-alcoholic fatty liver disease (NAFLD). Lipids in health and disease 2010, 9 (1), 1–9.

(3) Kesh, S. B.; Sarkar, D.; Manna, K. High-fat diet-induced oxidative stress and its impact on metabolic syndrome: a review. Asian J Pharm Clin Res 2016, 9 (1), 47–52.

(4) Kakehashi, A.; Stefanov, V. E.; Ishii, N.; Okuno, T.; Fujii, H.; Kawai, K.; Kawada, N.; Wanibuchi, H. Proteome Characteristics of Non-Alcoholic Steatohepatitis Liver Tissue and Associated Hepatocellular Carcinomas. 2017, 18 (2), 434.

(5) Iwaisako, K.; Brenner, D. A.; Kisseleva, T. What’s new in liver fibrosis? The origin of myofibroblasts in liver fibrosis. J Gastroenterol Hepatol 2012, 27 Suppl 2 (Suppl 2), 65-68. DOI: 10.1111/j.1440-1746.2011.07002.x

(6) Brunt, E. M. Pathology of nonalcoholic fatty liver disease. Nat Rev Gastroenterol Hepatol 2010, 7 (4), 195–203. DOI: 10.1038/nrgastro.2010.21

(7) Goldberg, D.; Ditah, I. C.; Saeian, K.; Lalehzari, M.; Aronsohn, A.; Gorospe, E. C.; Charlton, M. Changes in the prevalence of hepatitis C virus infection, nonalcoholic steatohepatitis, and alcoholic liver disease among patients with cirrhosis or liver failure on the waitlist for liver transplantation. Gastroenterology 2017, 152 (5), 1090–1099. e1091.

(8) Romeo, S.; Kozlitina, J.; Xing, C.; Pertsemlidis, A.; Cox, D.; Pennacchio, L. A.; Boerwinkle, E.; Cohen, J. C.; Hobbs, H. H. Genetic variation in PNPLA3 confers susceptibility to nonalcoholic fatty liver disease. Nature genetics 2008, 40 (12), 1461–1465.

(9) Tarantino, G.; Capone, D.; Finelli, C. Exposure to ambient air particulate matter and non-alcoholic fatty liver disease. World journal of gastroenterology: WJG 2013, 19 (25), 3951.

(10) Zhang, X.; Ji, X.; Wang, Q.; Li, J. Z. New insight into inter-organ crosstalk contributing to the pathogenesis of non-alcoholic fatty liver disease (NAFLD). Protein & cell 2018, 9 (2), 164–177.

(11) Fraulob, J. C.; Ogg-Diamantino, R.; Fernandes-Santos, C.; Aguila, M. B.; Mandarim-de-Lacerda, C. A. A mouse model of metabolic syndrome: insulin resistance, fatty liver and non-alcoholic fatty pancreas disease (NAFPD) in C57BL/6 mice fed a high fat diet. Journal of clinical biochemistry and nutrition 2010, 46 (3), 212–223.

(12) Chang, B. H.-J.; Li, L.; Saha, P.; Chan, L. Absence of adipose differentiation related protein upregulates hepatic VLDL secretion, relieves hepatosteatosis, and improves whole body insulin resistance in leptin-deficient mice [S]. Journal of lipid research 2010, 51 (8), 2132–2142.

(13) Spencer, M. D.; Hamp, T. J.; Reid, R. W.; Fischer, L. M.; Zeisel, S. H.; Fodor, A. A. Association between composition of the human gastrointestinal microbiome and development of fatty liver with choline deficiency. Gastroenterology 2011, 140 (3), 976–986.

(14) Al-Eryani, L.; Wahlang, B.; Falkner, K.; Guardiola, J.; Clair, H.; Prough, R.; Cave, M. Identification of environmental chemicals associated with the development of toxicant-associated fatty liver disease in rodents. Toxicologic pathology 2015, 43 (4), 482–497.

(15) Jain, R. B.; Ducatman, A. Selective associations of recent low concentrations of perfluoroalkyl substances with liver function biomarkers: NHANES 2011 to 2014 data on US adults aged≥ 20 years. Journal of Occupational and Environmental Medicine 2019, 61 (4), 293–302.

(16) Ortiz, L.; Nakamura, B.; Li, X.; Blumberg, B.; Luderer, U. Reprint of “In utero exposure to benzo [a] pyrene increases adiposity and causes hepatic steatosis in female mice, and glutathione deficiency is protective”. Toxicology letters 2014, 230 (2), 314–321.

(17) Vasiljevic, T.; Harner, T. Bisphenol A and its analogues in outdoor and indoor air: Properties, sources and global levels. Science of the Total Environment 2021, 789, 148013.

(18) Golub, M. S.; Wu, K. L.; Kaufman, F. L.; Li, L. H.; Moran■Messen, F.; Zeise, L.; Alexeeff, G. V.; Donald, J. M. Bisphenol A: developmental toxicity from early prenatal exposure a. Birth Defects Research Part B: Developmental and Reproductive Toxicology 2010, 89 (6), 441–466.

(19) Saili, K. S.; Corvi, M. M.; Weber, D. N.; Patel, A. U.; Das, S. R.; Przybyla, J.; Anderson, K. A.; Tanguay, R. L. Neurodevelopmental low-dose bisphenol A exposure leads to early life-stage hyperactivity and learning deficits in adult zebrafish. Toxicology 2012, 291 (1-3), 83–92.

(20) Wolstenholme, J. T.; Edwards, M.; Shetty, S. R.; Gatewood, J. D.; Taylor, J. A.; Rissman, E. F.; Connelly, J. J. Gestational exposure to bisphenol a produces transgenerational changes in behaviors and gene expression. Endocrinology 2012, 153 (8), 3828–3838.

(21) Bansal, A.; Li, C.; Xin, F.; Duemler, A.; Li, W.; Rashid, C.; Bartolomei, M. S.; Simmons, R. A. Transgenerational effects of maternal bisphenol: a exposure on offspring metabolic health. Journal of Developmental Origins of Health and Disease 2019, 10 (2), 164–175. DOI: 10.1017/S2040174418000764

(22) Marmugi, A.; Ducheix, S.; Lasserre, F.; Polizzi, A.; Paris, A.; Priymenko, N.; Bertrand-Michel, J.; Pineau, T.; Guillou, H.; Martin, P. G.; Mselli-Lakhal, L. Low doses of bisphenol A induce gene expression related to lipid synthesis and trigger triglyceride accumulation in adult mouse liver. Hepatology 2012, 55 (2), 395–407. DOI: 10.1002/hep.24685

(23) Cabaton, N. J.; Canlet, C.; Wadia, P. R.; Tremblay-Franco, M.; Gautier, R.; Molina, J.; Sonnenschein, C.; Cravedi, J. P.; Rubin, B. S.; Soto, A. M.; Zalko, D. Effects of low doses of bisphenol A on the metabolome of perinatally exposed CD-1 mice. Environ Health Perspect 2013, 121 (5), 586–593. DOI: 10.1289/ehp.1205588

(24) Nadal, A.; Alonso-Magdalena, P.; Soriano, S.; Quesada, I.; Ropero, A. B. The pancreatic beta-cell as a target of estrogens and xenoestrogens: Implications for blood glucose homeostasis and diabetes. Mol Cell Endocrinol 2009, 304 (1-2), 63–68. DOI: 10.1016/j.mce.2009.02.016

(25) Trasande, L.; Attina, T. M.; Blustein, J. Association Between Urinary Bisphenol A Concentration and Obesity Prevalence in Children and Adolescents. JAMA 2012, 308 (11), 1113–1121. DOI: 10.1001/2012.jama.11461 %J JAMA (acccessed 4/7/2023).

(26) Alonso-Magdalena, P.; Morimoto, S.; Ripoll, C.; Fuentes, E.; Nadal, A. The Estrogenic Effect of Bisphenol A Disrupts Pancreatic β-Cell Function In Vivo and Induces Insulin Resistance. Environmental Health Perspectives 2006, 114 (1), 106–112. DOI: 10.1289/ehp.8451 (acccessed 2023/04/07).

27. Hassan, Z. K.; Elobeid, M. A.; Virk, P.; Omer, S. A.; ElAmin, M.; Daghestani, M. H.; AlOlayan, E. M. J. O. m.; longevity, c. Bisphenol A induces hepatotoxicity through oxidative stress in rat model. 2012, 2012.

(28) Lama, S.; Vanacore, D.; Diano, N.; Nicolucci, C.; Errico, S.; Dallio, M.; Federico, A.; Loguercio, C.; Stiuso, P. Ameliorative effect of Silybin on bisphenol A induced oxidative stress, cell proliferation and steroid hormones oxidation in HepG2 cell cultures. Sci Rep 2019, 9 (1), 3228. DOI: 10.1038/s41598-019-40105-8

(29) Polyzos, S. A.; Kountouras, J.; Deretzi, G.; Zavos, C.; Mantzoros, C. S. The emerging role of endocrine disruptors in pathogenesis of insulin resistance: a concept implicating nonalcoholic fatty liver disease. Curr Mol Med 2012, 12 (1), 68–82. DOI: 10.2174/156652412798376161

(30) Labaronne, E.; Pinteur, C.; Vega, N.; Pesenti, S.; Julien, B.; Meugnier-Fouilloux, E.; Vidal, H.; Naville, D.; Le Magueresse-Battistoni, B. Low-dose pollutant mixture triggers metabolic disturbances in female mice leading to common and specific features as compared to a high-fat diet. J Nutr Biochem 2017, 45, 83–93. DOI: 10.1016/j.jnutbio.2017.04.001

(31) Jiang, Y.; Xia, W.; Zhu, Y.; Li, X.; Wang, D.; Liu, J.; Chang, H.; Li, G.; Xu, B.; Chen, X. Mitochondrial dysfunction in early life resulted from perinatal bisphenol A exposure contributes to hepatic steatosis in rat offspring. Toxicology letters 2014, 228 (2), 85–92.

32. Pirozzi, C.; Lama, A.; Annunziata, C.; Cavaliere, G.; Ruiz-Fernandez, C.; Monnolo, A.; Comella, F.; Gualillo, O.; Stornaiuolo, M.; Mollica, M. P.; et al. Oral Bisphenol A Worsens Liver Immune-Metabolic and Mitochondrial Dysfunction Induced by High-Fat Diet in Adult Mice: Cross-Talk between Oxidative Stress and Inflammasome Pathway. In Antioxidants, 2020; Vol. 9.

(33) Wang, K.; Zhao, Z.; Ji, W. Bisphenol A induces apoptosis, oxidative stress and inflammatory response in colon and liver of mice in a mitochondria-dependent manner. Biomedicine & Pharmacotherapy 2019, 117, 109182.

(34) Kautzky-Willer, A.; Harreiter, J.; Pacini, G. J. E. r. Sex and gender differences in risk, pathophysiology and complications of type 2 diabetes mellitus. 2016, 37 (3), 278–316.

(35) Peters, S. A.; Muntner, P.; Woodward, M. J. C. Sex differences in the prevalence of, and trends in, cardiovascular risk factors, treatment, and control in the United States, 2001 to 2016. 2019, 139 (8), 1025-1035.

(36) DeBenedictis, B.; Guan, H.; Yang, K. J. J. o. C. B. Prenatal exposure to bisphenol A disrupts mouse fetal liver maturation in a sex■specific manner. 2016, 117 (2), 344–350.

(37) Strakovsky, R. S.; Wang, H.; Engeseth, N. J.; Flaws, J. A.; Helferich, W. G.; Pan, Y.-X.; Lezmi, S. Developmental bisphenol A (BPA) exposure leads to sex-specific modification of hepatic gene expression and epigenome at birth that may exacerbate high-fat diet-induced hepatic steatosis. Toxicology and Applied Pharmacology 2015, 284 (2), 101–112. DOI: 10.1016/j.taap.2015.02.021

(38) Hijazi, A.; Guan, H.; Cernea, M.; Yang, K. J. T. F. J. Prenatal exposure to bisphenol A disrupts mouse fetal lung development. 2015, 29 (12), 4968–4977.

(39) Tainaka, H.; Takahashi, H.; Umezawa, M.; Tanaka, H.; Nishimune, Y.; Oshio, S.; Takeda, K. J. T. J. o. t. s. Evaluation of the testicular toxicity of prenatal exposure to bisphenol A based on microarray analysis combined with MeSH annotation. 2012, 37 (3), 539–548.

(40) Lombó, M.; Fernández-Díez, C.; González-Rojo, S.; Herráez, M. P. J. S. R. Genetic and epigenetic alterations induced by bisphenol A exposure during different periods of spermatogenesis: from spermatozoa to the progeny. 2019, 9 (1), 1–13.

(41) Manikkam, M.; Tracey, R.; Guerrero-Bosagna, C.; Skinner, M. K. J. P. o. Plastics derived endocrine disruptors (BPA, DEHP and DBP) induce epigenetic transgenerational inheritance of obesity, reproductive disease and sperm epimutations. 2013, 8 (1), e55387.

(42) Zheng, H.; Zhou, X.; Li, D.-k.; Yang, F.; Pan, H.; Li, T.; Miao, M.; Li, R.; Yuan, W. J. P. O. Genome-wide alteration in DNA hydroxymethylation in the sperm from bisphenol A-exposed men. 2017, 12 (6), e0178535.

(43) Matsumoto, T.; Terai, S.; Oishi, T.; Kuwashiro, S.; Fujisawa, K.; Yamamoto, N.; Fujita, Y.; Hamamoto, Y.; Furutani-Seiki, M.; Nishina, H. Medaka as a model for human nonalcoholic steatohepatitis. Disease models & mechanisms 2010, 3 (7-8), 431–440.

(44) Chakraborty, S.; Dissanayake, M.; Godwin, J.; Wang, X.; Bhandari, R. K. Ancestral BPA exposure caused defects in the liver of medaka for four generations. Science of The Total Environment 2023, 856, 159067. DOI: 10.1016/j.scitotenv.2022.159067

(45) Matsumoto, T.; Terai, S.; Oishi, T.; Kuwashiro, S.; Fujisawa, K.; Yamamoto, N.; Fujita, Y.; Hamamoto, Y.; Furutani-Seiki, M.; Nishina, H.; Sakaida, I. Medaka as a model for human nonalcoholic steatohepatitis. Disease Models & Mechanisms 2010, 3 (7-8), 431–440. DOI: 10.1242/dmm.002311 %J Disease Models & Mechanisms (acccessed 4/13/2023).

(46) Ryan, K. K.; Haller, A. M.; Sorrell, J. E.; Woods, S. C.; Jandacek, R. J.; Seeley, R. J. Perinatal exposure to bisphenol-a and the development of metabolic syndrome in CD-1 mice. Endocrinology 2010, 151 (6), 2603–2612. DOI: 10.1210/en.2009-1218

(47) Angle, B. M.; Do, R. P.; Ponzi, D.; Stahlhut, R. W.; Drury, B. E.; Nagel, S. C.; Welshons, W. V.; Besch-Williford, C. L.; Palanza, P.; Parmigiani, S.;, et al. Metabolic disruption in male mice due to fetal exposure to low but not high doses of bisphenol A (BPA): evidence for effects on body weight, food intake, adipocytes, leptin, adiponectin, insulin and glucose regulation. Reprod Toxicol 2013, 42, 256–268. DOI: 10.1016/j.reprotox.2013.07.017

(48) Puttabyatappa, M.; Martin, J. D.; Andriessen, V.; Stevenson, M.; Zeng, L.; Pennathur, S.; Padmanabhan, V. Developmental programming: Changes in mediators of insulin sensitivity in prenatal bisphenol A-treated female sheep. Reprod Toxicol 2019, 85, 110–122. DOI: 10.1016/j.reprotox.2019.03.002

(49) Samardzija, D.; Pogrmic-Majkic, K.; Fa, S.; Stanic, B.; Jasnic, J.; Andric, N. Bisphenol A decreases progesterone synthesis by disrupting cholesterol homeostasis in rat granulosa cells. Mol Cell Endocrinol 2018, 461, 55–63. DOI: 10.1016/j.mce.2017.08.013

(50) Yamamoto, T.; Yasuhara, A.; Shiraishi, H.; Nakasugi, O. Bisphenol A in hazardous waste landfill leachates. Chemosphere 2001, 42 (4), 415–418.

(51) Crain, D. A.; Eriksen, M.; Iguchi, T.; Jobling, S.; Laufer, H.; LeBlanc, G. A.; Guillette Jr, L. J. An ecological assessment of bisphenol-A: evidence from comparative biology. Reproductive toxicology 2007, 24 (2), 225–239.

(52) Thayil, A. J.; Wang, X.; Bhandari, P.; vom Saal, F. S.; Tillitt, D. E.; Bhandari, R. K. Bisphenol A and 17α-ethinylestradiol-induced transgenerational gene expression differences in the brain– pituitary–testis axis of medaka, Oryzias latipes†. Biology of Reproduction 2020, 103 (6), 1324–1335. DOI: 10.1093/biolre/ioaa169 (acccessed 11/26/2023).

(53) Bhandari, R. K.; Wang, X.; Vom Saal, F. S.; Tillitt, D. E. Transcriptome analysis of testis reveals the effects of developmental exposure to bisphenol a or 17α-ethinylestradiol in medaka (Oryzias latipes). Aquatic Toxicology 2020, 225, 105553.

(54) Wang, X.; Bhandari, R. K. The dynamics of DNA methylation during epigenetic reprogramming of primordial germ cells in medaka (Oryzias latipes). Epigenetics 2020, 15 (5), 483–498.

(55) Van Wettere, A. J. Pathogenesis of Liver Fibrosis and Regeneration in the Japanese Medaka (Oryzias latipes); North Carolina State University, 2012.

(56) Wang, X.; Bhandari, R. K. DNA methylation dynamics during epigenetic reprogramming of medaka embryo. Epigenetics 2019, 14 (6), 611–622.

(57) Bhandari, R. K.; Vom Saal, F. S.; Tillitt, D. E. Transgenerational effects from early developmental exposures to bisphenol A or 17α-ethinylestradiol in medaka, Oryzias latipes. Scientific reports 2015, 5 (1), 9303.

(58) Wang, X.; Hill, D.; Tillitt, D. E.; Bhandari, R. K. Bisphenol A and 17alpha-ethinylestradiol-induced transgenerational differences in expression of osmoregulatory genes in the gill of medaka (Oryzias latipes). Aquat Toxicol 2019, 211, 227–234. DOI: 10.1016/j.aquatox.2019.04.005

(59) Thayil, A. J.; Wang, X.; Bhandari, P.; Vom Saal, F. S.; Tillitt, D. E.; Bhandari, R. K. Bisphenol A and 17α-ethinylestradiol-induced transgenerational gene expression differences in the brain-pituitary-testis axis of medaka, Oryzias latipes†. Biol Reprod 2020, 103 (6), 1324–1335. DOI: 10.1093/biolre/ioaa169

(60) Wang, X.; Hill, D.; Tillitt, D. E.; Bhandari, R. K. Bisphenol A and 17α-ethinylestradiol-induced transgenerational differences in expression of osmoregulatory genes in the gill of medaka (Oryzias latipes). Aquat Toxicol 2019, 211, 227–234. DOI: 10.1016/j.aquatox.2019.04.005

(61) Aranda, P. S.; LaJoie, D. M.; Jorcyk, C. L. Bleach gel: a simple agarose gel for analyzing RNA quality. Electrophoresis 2012, 33 (2), 366–369.

(62) Chen, S.; Zhou, Y.; Chen, Y.; Gu, J. fastp: an ultra-fast all-in-one FASTQ preprocessor. Bioinformatics 2018, 34 (17), i884–i890. DOI: 10.1093/bioinformatics/bty560 (acccessed 12/18/2022).

(63) Dobin, A.; Davis, C. A.; Schlesinger, F.; Drenkow, J.; Zaleski, C.; Jha, S.; Batut, P.; Chaisson, M.; Gingeras, T. R. STAR: ultrafast universal RNA-seq aligner. Bioinformatics 2013, 29 (1), 15–21. DOI: 10.1093/bioinformatics/bts635 (acccessed 12/18/2022).

(64) Love, M. I.; Anders, S.; Huber, W. Analyzing RNA-seq data with DESeq2. Bioconductor 2017, 2, 1–63.

(65) Hammad, A.; Zheng, Z.-H.; Namani, A.; Elshaer, M.; Wang, X. J.; Tang, X. Transcriptome analysis of potential candidate genes and molecular pathways in colitis-associated colorectal cancer of Mkp-1-deficient mice. BMC cancer 2021, 21 (1), 1–13.

(66) Lemecha, M.; Chalise, J. P.; Takamuku, Y.; Zhang, G.; Yamakawa, T.; Larson, G.; Itakura, K. Lcn2 mediates adipocyte-muscle-tumor communication and hypothermia in pancreatic cancer cachexia. Molecular Metabolism 2022, 66, 101612.

(67) Luna, A.; Babur, Ö.; Aksoy, B. A.; Demir, E.; Sander, C. PaxtoolsR: pathway analysis in R using Pathway Commons. Bioinformatics 2016, 32 (8), 1262–1264.

(68) Nguyen, H. T.; Yamamoto, K.; Iida, M.; Agusa, T.; Ochiai, M.; Guo, J.; Karthikraj, R.; Kannan, K.; Kim, E.-Y.; Iwata, H. Effects of prenatal bisphenol A exposure on the hepatic transcriptome and proteome in rat offspring. Science of The Total Environment 2020, 720, 137568. DOI: 10.1016/j.scitotenv.2020.137568

(69) Greco, D.; Kotronen, A.; Westerbacka, J.; Puig, O.; Arkkila, P.; Kiviluoto, T.; Laitinen, S.; Kolak, M.; Fisher, R. M.; Hamsten, A.; et al. Gene expression in human NAFLD. 2008, 294 (5), G1281–G1287. DOI: 10.1152/ajpgi.00074.2008

(70) Teufel, A.; Itzel, T.; Erhart, W.; Brosch, M.; Wang, X. Y.; Kim, Y. O.; von Schönfels, W.; Herrmann, A.; Brückner, S.; Stickel, F.;, et al. Comparison of Gene Expression Patterns Between Mouse Models&#xa0;of Nonalcoholic Fatty Liver Disease and Liver Tissues&#xa0;From Patients. Gastroenterology 2016, 151 (3), 513–525.e510. DOI: 10.1053/j.gastro.2016.05.051 (acccessed 2023/11/30).

(71) Diamante, G.; Cely, I.; Zamora, Z.; Ding, J.; Blencowe, M.; Lang, J.; Bline, A.; Singh, M.; Lusis, A. J.; Yang, X. Systems toxicogenomics of prenatal low-dose BPA exposure on liver metabolic pathways, gut microbiota, and metabolic health in mice. Environment International 2021, 146, 106260. DOI: 10.1016/j.envint.2020.106260

(72) Fan, J.; Yang, M.; Liang, C.; Liang, C.; Guo, J. The Diagnostic and Prognostic Value of BEN-Domain (BEND) Family Genes in Gastric Cancer. 2022.

(73) Ding, Z.; Ericksen, R. E.; Escande-Beillard, N.; Lee, Q. Y.; Loh, A.; Denil, S.; Steckel, M.; Haegebarth, A.; Ho, T. S. W.; Chow, P. J. J. o. h. Metabolic pathway analyses identify proline biosynthesis pathway as a promoter of liver tumorigenesis. 2020, 72 (4), 725–735.

(74) Warnier, M.; Roudbaraki, M.; Derouiche, S.; Delcourt, P.; Bokhobza, A.; Prevarskaya, N.; Mariot, P. J. O. CACNA2D2 promotes tumorigenesis by stimulating cell proliferation and angiogenesis. 2015, 34 (42), 5383–5394.

(75) Wang, S.; Gu, L.; Huang, L.; Fang, J.; Liu, Z.; Xu, Q. J. B.; Communications, B. R. The upregulation of PYCR2 is associated with aggressive colon cancer progression and a poor prognosis. 2021, 572, 20–26.

(76) Ding, Z.; Ericksen, R. E.; Escande-Beillard, N.; Lee, Q. Y.; Loh, A.; Denil, S.; Steckel, M.; Haegebarth, A.; Wai Ho, T. S.; Chow, P.;, et al. Metabolic pathway analyses identify proline biosynthesis pathway as a promoter of liver tumorigenesis. Journal of Hepatology 2020, 72 (4), 725–735. DOI: 10.1016/j.jhep.2019.10.026

(77) Lee, A.-H.; Scapa, E. F.; Cohen, D. E.; Glimcher, L. H. J. S. Regulation of hepatic lipogenesis by the transcription factor XBP1. 2008, *320* (5882), 1492–1496.

(78) Gonzalez-Rodriguez, A.; Mayoral, R.; Agra, N.; Valdecantos, M.; Pardo, V.; Miquilena-Colina, M.; Vargas-Castrillón, J.; Lo Iacono, O.; Corazzari, M.; Fimia, G. J. C. d.; disease. Impaired autophagic flux is associated with increased endoplasmic reticulum stress during the development of NAFLD. 2014, 5 (4), e1179–e1179.

(79) Gardner, B. M.; Walter, P. J. S. Unfolded proteins are Ire1-activating ligands that directly induce the unfolded protein response. 2011, 333 (6051), 1891–1894.

(80) Lee, A.-H.; Scapa, E. F.; Cohen, D. E.; Glimcher, L. H. Regulation of Hepatic Lipogenesis by the Transcription Factor XBP1. 2008, 320 (5882), 1492–1496. DOI: doi:10.1126/science.1158042

(81) Li, K.; Xiao, Y.; Yu, J.; Xia, T.; Liu, B.; Guo, Y.; Deng, J.; Chen, S.; Wang, C.; Guo, F. J. J. o. B. C. Liver-specific gene inactivation of the transcription factor ATF4 alleviates alcoholic liver steatosis in mice. 2016, 291 (35), 18536–18546.

(82) Xiao, W.; Wang, J.; Ou, C.; Zhang, Y.; Ma, L.; Weng, W.; Pan, Q.; Sun, F. J. B.; communications, b. r. Mutual interaction between YAP and c-Myc is critical for carcinogenesis in liver cancer. 2013, 439 (2), 167–172.

(83) Yang, T.; Poenisch, M.; Khanal, R.; Hu, Q.; Dai, Z.; Li, R.; Song, G.; Yuan, Q.; Yao, Q.; Shen, X.;, et al. Therapeutic HNF4A mRNA attenuates liver fibrosis in a preclinical model. Journal of Hepatology 2021, 75 (6), 1420–1433. DOI: 10.1016/j.jhep.2021.08.011

(84) Ke, Z.-H.; Pan, J.-X.; Jin, L.-Y.; Xu, H.-Y.; Yu, T.-T.; Ullah, K.; Rahman, T. U.; Ren, J.; Cheng, Y.; Dong, X.-Y. J. S. r. Bisphenol A exposure may induce hepatic lipid accumulation via reprogramming the DNA methylation patterns of genes involved in lipid metabolism. 2016, 6 (1), 31331.

(85) Gu, Z.; Jia, R.; He, Q.; Cao, L.; Du, J.; Feng, W.; Jeney, G.; Xu, P.; Yin, G. Alteration of lipid metabolism, autophagy, apoptosis and immune response in the liver of common carp (Cyprinus carpio) after long-term exposure to bisphenol A. Ecotoxicology and Environmental Safety 2021, 211, 111923. DOI: 10.1016/j.ecoenv.2021.111923

(86) Hong, L.; Xu, Y.; Wang, D.; Zhang, Q.; Li, X.; Xie, C.; Wu, J.; Zhong, C.; Fu, J.; Geng, S. Sulforaphane ameliorates bisphenol A-induced hepatic lipid accumulation by inhibiting endoplasmic reticulum stress. Scientific Reports 2023, 13 (1), 1147. DOI: 10.1038/s41598-023-28395-5

(87) Eferl, R.; Casanova, E.; Österreicher, C. H.; Blaas, L.; Mair, M. J. F. i. B.-L. JAK-STAT signaling in hepatic fibrosis. 2011, 16 (8), 2794–2811.

(88) Biernacka, A.; Dobaczewski, M.; Frangogiannis, N. G. J. G. f. TGF-β signaling in fibrosis. 2011, 29 (5), 196–202.

(89) Guzman, E. Transgenerational differences in gut microbiota population and epigenetic responses of host intestinal epithelial cells; The University of North Carolina at Greensboro, 2021.

(90) Tan, M.; Ye, J.; Zhao, M.; Ke, X.; Huang, K.; Liu, H. Recent developments in the regulation of cholesterol transport by natural molecules. 2021, 35 (10), 5623–5633. DOI: 10.1002/ptr.7198

(91) Ouimet, M.; Barrett, T. J.; Fisher, E. A. J. C. r. HDL and reverse cholesterol transport: Basic mechanisms and their roles in vascular health and disease. 2019, 124 (10), 1505–1518.

(92) Choi, S.; Fong, L.; Kirven, M.; Cooper, A. J. T. J. o. c. i. Use of an anti-low density lipoprotein receptor antibody to quantify the role of the LDL receptor in the removal of chylomicron remnants in the mouse in vivo. 1991, 88 (4), 1173–1181.

(93) Bieghs, V.; Van Gorp, P. J.; Wouters, K.; Hendrikx, T.; Gijbels, M. J.; van Bilsen, M.; Bakker, J.; Binder, C. J.; Lütjohann, D.; Staels, B.;, et al. LDL Receptor Knock-Out Mice Are a Physiological Model Particularly Vulnerable to Study the Onset of Inflammation in Non-Alcoholic Fatty Liver Disease. PLOS ONE 2012, 7 (1), e30668. DOI: 10.1371/journal.pone.0030668

(94) Somm, E.; Schwitzgebel, V. M.; Toulotte, A.; Cederroth, C. R.; Combescure, C.; Nef, S.; Aubert, M. L.; Hüppi, P. S. Perinatal Exposure to Bisphenol A Alters Early Adipogenesis in the Rat. 2009, 117 (10), 1549–1555. DOI: doi:10.1289/ehp.11342

(95) Vega□Badillo, J.; Gutiérrez□Vidal, R.; Hernández□Pérez, H. A.; Villamil□Ramírez, H.; León□Mimila, P.; Sánchez□Muñoz, F.; Morán□Ramos, S.; Larrieta□Carrasco, E.; Fernández□Silva, I.; Méndez□Sánchez, N. J. L. I. Hepatic miR□33a/miR□144 and their target gene ABCA1 are associated with steatohepatitis in morbidly obese subjects. 2016, 36 (9), 1383–1391.

(96) Miyazaki, M.; Dobrzyn, A.; Man, W. C.; Chu, K.; Sampath, H.; Kim, H.-J.; Ntambi, J. M. J. J. o. b. c. Stearoyl-CoA desaturase 1 gene expression is necessary for fructose-mediated induction of lipogenic gene expression by sterol regulatory element-binding protein-1c-dependent and-independent mechanisms. 2004, 279 (24), 25164–25171.

(97) Jiang, G.; Li, Z.; Liu, F.; Ellsworth, K.; Dallas-Yang, Q.; Wu, M.; Ronan, J.; Esau, C.; Murphy, C.; Szalkowski, D. J. T. J. o. c. i. Prevention of obesity in mice by antisense oligonucleotide inhibitors of stearoyl-CoA desaturase–1. 2005, 115 (4), 1030–1038.

(98) Ip, E.; Farrell, G. C.; Robertson, G.; Hall, P.; Kirsch, R.; Leclercq, I. Central role of PPARα-dependent hepatic lipid turnover in dietary steatohepatitis in mice. 2003, 38 (1), 123–132. DOI: 10.1053/jhep.2003.50307

(99) Morán-Salvador, E.; Titos, E.; Rius, B.; González-Périz, A.; García-Alonso, V.; López-Vicario, C.; Miquel, R.; Barak, Y.; Arroyo, V.; Clària, J. Cell-specific PPAR&#x3b3; deficiency establishes anti-inflammatory and anti-fibrogenic properties for this nuclear receptor in non-parenchymal liver cells. Journal of Hepatology 2013, 59 (5), 1045–1053. DOI: 10.1016/j.jhep.2013.06.023 (acccessed 2023/05/13).

(100) Schreurs, M.; Kuipers, F.; Van Der Leij, F. R. Regulatory enzymes of mitochondrial β-oxidation as targets for treatment of the metabolic syndrome. 2010, 11 (5), 380–388. DOI: 10.1111/j.1467-789X.2009.00642.x

(101) Strakovsky, R. S.; Wang, H.; Engeseth, N. J.; Flaws, J. A.; Helferich, W. G.; Pan, Y.-X.; Lezmi, S. J. T.; pharmacology, a. Developmental bisphenol A (BPA) exposure leads to sex-specific modification of hepatic gene expression and epigenome at birth that may exacerbate high-fat diet-induced hepatic steatosis. 2015, 284 (2), 101–112.

(102) Brown, Z. J.; Fu, Q.; Ma, C.; Kruhlak, M.; Zhang, H.; Luo, J.; Heinrich, B.; Yu, S. J.; Zhang, Q.; Wilson, A.;, et al. Carnitine palmitoyltransferase gene upregulation by linoleic acid induces CD4(+) T cell apoptosis promoting HCC development. Cell Death Dis 2018, 9 (6), 620. DOI: 10.1038/s41419-018-0687-6

(103) Ke, Z.-H.; Pan, J.-X.; Jin, L.-Y.; Xu, H.-Y.; Yu, T.-T.; Ullah, K.; Rahman, T. U.; Ren, J.; Cheng, Y.; Dong, X.-Y.;, et al. Bisphenol A Exposure May Induce Hepatic Lipid Accumulation via Reprogramming the DNA Methylation Patterns of Genes Involved in Lipid Metabolism. Scientific Reports 2016, 6 (1), 31331. DOI: 10.1038/srep31331

(104) Dolt, K. S.; Karar, J.; Mishra, M. K.; Salim, J.; Kumar, R.; Grover, S. K.; Qadar Pasha, M. A. Transcriptional downregulation of sterol metabolism genes in murine liver exposed to acute hypobaric hypoxia. Biochemical and Biophysical Research Communications 2007, 354 (1), 148–153. DOI: 10.1016/j.bbrc.2006.12.159

(105) Miquilena-Colina, M. E.; Lima-Cabello, E.; Sánchez-Campos, S.; García-Mediavilla, M. V.; Fernández-Bermejo, M.; Lozano-Rodríguez, T.; Vargas-Castrillón, J.; Buqué, X.; Ochoa, B.; Aspichueta, P. J. G. Hepatic fatty acid translocase CD36 upregulation is associated with insulin resistance, hyperinsulinaemia and increased steatosis in non-alcoholic steatohepatitis and chronic hepatitis C. 2011, 60 (10), 1394–1402.

(106) Rada, P.; González-Rodríguez, Á.; García-Monzón, C.; Valverde, Á. M. J. C. d.; disease. Understanding lipotoxicity in NAFLD pathogenesis: is CD36 a key driver? 2020, 11 (9), 802.

(107) Bechmann, L. P.; Gieseler, R. K.; Sowa, J. P.; Kahraman, A.; Erhard, J.; Wedemeyer, I.; Emons, B.; Jochum, C.; Feldkamp, T.; Gerken, G. J. L. I. Apoptosis is associated with CD36/fatty acid translocase upregulation in non□alcoholic steatohepatitis. 2010, 30 (6), 850–859.

(108) Pacana, T.; Cazanave, S.; Verdianelli, A.; Patel, V.; Min, H.-K.; Mirshahi, F.; Quinlivan, E.; Sanyal, A. J. J. P. o. Dysregulated hepatic methionine metabolism drives homocysteine elevation in diet-induced nonalcoholic fatty liver disease. 2015, 10 (8), e0136822.

(109) Kremer, M.; Hines, I. N.; Milton, R. J.; Wheeler, M. D. J. H. Favored T helper 1 response in a mouse model of hepatosteatosis is associated with enhanced T cell–mediated hepatitis. 2006, 44 (1), 216–227.

(110) Bertola, A.; Bonnafous, S.; Anty, R.; Patouraux, S.; Saint-Paul, M.-C.; Iannelli, A.; Gugenheim, J.; Barr, J.; Mato, J. M.; Le Marchand-Brustel, Y. J. P. o. Hepatic expression patterns of inflammatory and immune response genes associated with obesity and NASH in morbidly obese patients. 2010, 5 (10), e13577.

(111) Kim, S.; Mun, G.-i.; Choi, E.; Kim, M.; Jeong, J. S.; Kang, K. W.; Jee, S.; Lim, K.-M.; Lee, Y.-S. Submicromolar bisphenol A induces proliferation and DNA damage in human hepatocyte cell lines in vitro and in juvenile rats in vivo. Food and Chemical Toxicology 2018, 111, 125–132. DOI: 10.1016/j.fct.2017.11.010

(112) Li, Z.; Wang, F.; Liang, B.; Su, Y.; Sun, S.; Xia, S.; Shao, J.; Zhang, Z.; Hong, M.; Zhang, F. J. S. t.; therapy, t. Methionine metabolism in chronic liver diseases: an update on molecular mechanism and therapeutic implication. 2020, 5 (1), 280.

(113) Pfeifer, D.; Chung, Y. M.; Hu, M. C. J. E. h. p. Effects of low-dose bisphenol A on DNA damage and proliferation of breast cells: the role of c-Myc. 2015, 123 (12), 1271–1279.

(114) Bindhumol, V.; Chitra, K.; Mathur, P. J. T. Bisphenol A induces reactive oxygen species generation in the liver of male rats. 2003, 188 (2-3), 117–124.

(115) Izzotti, A.; Kanitz, S.; D’Agostini, F.; Camoirano, A.; De Flora, S. J. M. R. G. T.; Mutagenesis, E. Formation of adducts by bisphenol A, an endocrine disruptor, in DNA in vitro and in liver and mammary tissue of mice. 2009, 679 (1-2), 28–32.

(116) Rolo, A. P.; Teodoro, J. S.; Palmeira, C. M. J. F. r. b.; medicine. Role of oxidative stress in the pathogenesis of nonalcoholic steatohepatitis. 2012, 52 (1), 59–69.

(117) Mantena, S. K.; King, A. L.; Andringa, K. K.; Eccleston, H. B.; Bailey, S. M. J. F. R. B.; Medicine. Mitochondrial dysfunction and oxidative stress in the pathogenesis of alcohol-and obesity-induced fatty liver diseases. 2008, 44 (7), 1259–1272.

(118) Glimcher, L. H.; Lee, A. H. J. A. o. t. N. Y. A. o. S. From sugar to fat: How the transcription factor XBP1 regulates hepatic lipogenesis. 2009, 1173, E2–E9.

(119) Hotamisligil, G. S. J. C. Endoplasmic reticulum stress and the inflammatory basis of metabolic disease. 2010, 140 (6), 900–917.

(120) Kim, I.; Xu, W.; Reed, J. C. J. N. r. D. d. Cell death and endoplasmic reticulum stress: disease relevance and therapeutic opportunities. 2008, 7 (12), 1013–1030.

(121) Lee, C.-W.; Wong, L. L.-Y.; Tse, E. Y.-T.; Liu, H.-F.; Leong, V. Y.-L.; Lee, J. M.-F.; Hardie, D. G.; Ng, I. O.-L.; Ching, Y.-P. J. C. r. AMPK promotes p53 acetylation via phosphorylation and inactivation of SIRT1 in liver cancer cells. 2012, 72 (17), 4394–4404.

(122) Ferrín, G.; Guerrero, M.; Amado, V.; Rodríguez-Perálvarez, M.; De la Mata, M. J. I. J. o. M. S. Activation of mTOR signaling pathway in hepatocellular carcinoma. 2020, 21 (4), 1266.

(123) Han, J.; Wang, Y. J. P.; cell. mTORC1 signaling in hepatic lipid metabolism. 2018, 9 (2), 145–151.

(124) Bailey, C. L.; Kelly, P.; Casey, P. J. J. C. r. Activation of Rap1 promotes prostate cancer metastasis. 2009, 69 (12), 4962–4968.

(125) Uno, S.; Nebert, D. W.; Makishima, M. Cytochrome P450 1A1 (CYP1A1) protects against nonalcoholic fatty liver disease caused by Western diet containing benzo[a]pyrene in mice. Food and Chemical Toxicology 2018, 113, 73–82. DOI: 10.1016/j.fct.2018.01.029

(126) Uno, S.; Dalton, T. P.; Sinclair, P. R.; Gorman, N.; Wang, B.; Smith, A. G.; Miller, M. L.; Shertzer, H. G.; Nebert, D. W. Cyp1a1(−/−) male mice: protection against high-dose TCDD-induced lethality and wasting syndrome, and resistance to intrahepatocyte lipid accumulation and uroporphyria. Toxicology and Applied Pharmacology 2004, 196 (3), 410–421. DOI: 10.1016/j.taap.2004.01.014

(127) Zhang, Y.; Li, F.; Patterson, A. D.; Wang, Y.; Krausz, K. W.; Neale, G.; Thomas, S.; Nachagari, D.; Vogel, P.; Vore, M. Abcb11 deficiency induces cholestasis coupled to impaired β-fatty acid oxidation in mice. Journal of Biological Chemistry 2012, 287 (29), 24784–24794.

(128) Dichtel, L. E.; Corey, K. E.; Misdraji, J.; Bredella, M. A.; Schorr, M.; Osganian, S. A.; Young, B. J.; Sung, J. C.; Miller, K. K. The Association Between IGF-1 Levels and the Histologic Severity of Nonalcoholic Fatty Liver Disease. Clin Transl Gastroenterol 2017, 8 (1), e217. DOI: 10.1038/ctg.2016.72

(129) Shi, Z.; Zhang, K.; Chen, T.; Zhang, Y.; Du, X.; Zhao, Y.; Shao, S.; Zheng, L.; Han, T.; Hong, W. Transcriptional factor ATF3 promotes liver fibrosis via activating hepatic stellate cells. Cell Death & Disease 2020, 11 (12), 1066. DOI: 10.1038/s41419-020-03271-6

(130) Inaba, Y.; Hashiuchi, E.; Watanabe, H.; Kimura, K.; Oshima, Y.; Tsuchiya, K.; Murai, S.; Takahashi, C.; Matsumoto, M.; Kitajima, S.;, et al. The transcription factor ATF3 switches cell death from apoptosis to necroptosis in hepatic steatosis in male mice. Nature Communications 2023, 14 (1), 167. DOI: 10.1038/s41467-023-35804-w

(131) Ohata, T.; Yokoo, H.; Kamiyama, T.; Fukai, M.; Aiyama, T.; Hatanaka, Y.; Hatanaka, K.; Wakayama, K.; Orimo, T.; Kakisaka, T.;, et al. Fatty acid-binding protein 5 function in hepatocellular carcinoma through induction of epithelial–mesenchymal transition. Cancer Medicine 2017, 6 (5), 1049–1061. DOI: 10.1002/cam4.1020 (acccessed 2023/11/29).

(132) Liu, J.-Y.; Chiang, T.; Liu, C.-H.; Chern, G.-G.; Lin, T.-T.; Gao, D.-Y.; Chen, Y. Delivery of siRNA using CXCR4-targeted nanoparticles modulates tumor microenvironment and achieves a potent antitumor response in liver cancer. Molecular Therapy 2015, 23 (11), 1772–1782.

(133) Liu, H.; Liu, Y.; Liu, W.; Zhang, W.; Xu, J. EZH2-mediated loss of miR-622 determines CXCR4 activation in hepatocellular carcinoma. Nature communications 2015, 6 (1), 8494.

134. Li, S.; Raza, S. H.; Zhao, C.; Cheng, G.; Zan, L. Overexpression of PLIN1 Promotes Lipid Metabolism in Bovine Adipocytes. In Animals, 2020; Vol. 10.

(135) Kammel, A.; Saussenthaler, S.; Jähnert, M.; Jonas, W.; Stirm, L.; Hoeflich, A.; Staiger, H.; Fritsche, A.; Häring, H.-U.; Joost, H.-G. J. H. m. g. Early hypermethylation of hepatic Igfbp2 results in its reduced expression preceding fatty liver in mice. 2016, 25 (12), 2588–2599.

(136) Jiang, J.; Briedé, J. J.; Jennen, D. G. J.; Van Summeren, A.; Saritas-Brauers, K.; Schaart, G.; Kleinjans, J. C. S.; de Kok, T. M. C. M. Increased mitochondrial ROS formation by acetaminophen in human hepatic cells is associated with gene expression changes suggesting disruption of the mitochondrial electron transport chain. Toxicology Letters 2015, 234 (2), 139–150. DOI: 10.1016/j.toxlet.2015.02.012

(137) Wruck, W.; Adjaye, J. Meta-analysis reveals up-regulation of cholesterol processes in non-alcoholic and down-regulation in alcoholic fatty liver disease. World J Hepatol 2017, 9 (8), 443–454. DOI: 10.4254/wjh.v9.i8.443

(138) Clemmons, D. R. J. T. J. o. c. i. The relative roles of growth hormone and IGF-1 in controlling insulin sensitivity. 2004, 113 (1), 25–27.

(139) García-Galiano, D.; Sánchez-Garrido, M. A.; Espejo, I.; Montero, J. L.; Costán, G.; Marchal, T.; Membrives, A.; Gallardo-Valverde, J. M.; Muñoz-Castañeda, J. R.; Arévalo, E. J. O. s. IL-6 and IGF-1 are independent prognostic factors of liver steatosis and non-alcoholic steatohepatitis in morbidly obese patients. 2007, 17, 493–503.

